# A chemical tool for improved culture of human pluripotent stem cells

**DOI:** 10.1101/2020.01.24.918268

**Authors:** Laurence Silpa, Maximilian Schuessler, Gu Liu, Marcus Olivecrona, Lucia Groizard-Payeras, Elizabeth Couper, Carole J. R. Bataille, Mark Stevenson, Len W. Seymour, Stephen G. Davies, William S. James, Sally A. Cowley, Angela J. Russell

## Abstract

The large-scale and cost-effective production of quality-controlled human pluripotent stem cells (hPSC) for use in cell therapy and drug discovery requires chemically-defined xenobiotic-free culture systems that enable easy and homogeneous expansion of pluripotent cells. Through phenotypic screening, we have identified a small molecule, OXS8360 (an optimized derivative of (-)-Indolactam V ((-)-ILV)), that stably disrupts hPSC cell-cell contacts. Proliferation of hPSC in OXS8360 is normal, as are pluripotency signatures, directed differentiation to hallmark lineages and karyotype over extended passaging. In 3D culture, OXS8360-treated hPSC form smaller, more uniform aggregates, that are easier to dissociate, greatly facilitating expansion. The mode of action of OXS8360 involves disruption of the localisation of the cell-cell adhesion molecule E-cadherin, via activation of unconventional Protein Kinase C isoforms. OXS8360 media supplementation is therefore able to yield more uniform, disaggregated 2D and 3D hPSC cultures, providing the hPSC field with an affordable tool to improve hPSC quality and scalability.

## Introduction

Human pluripotent stem cells (hPSC) have the unique properties of indefinite self-renewal and the ability to differentiate into representatives of the three germ layers^1^. However, a fundamental requirement needed to exploit the full potential of hPSC, in both research and medicine, is the ability to produce large numbers of cells of consistent quality and in a cost effective manner. In addition, the application of hPSC also requires ease of use, and good manufacturing practice (GMP) compatibility, without compromising pluripotency or increasing heterogeneity in the culture^2^. While hPSC are most commonly grown in chemically defined media, such as mTeSR1™ ^3^ or E8^4^, the main challenge encountered in traditional 2D culture methods is the limited quantities in which hPSC can be produced for experimental and therapeutic application. Zweigerdt *et al.*^5^ have shown that at least 2×10^9^ differentiated cells of the representative lineage would be necessary to treat one individual in the case of heart repair or β-cell replacement in type 1 diabetes, which requires about 500 densely grown 10-cm dishes and complex automation. One strategy for overcoming these difficulties is to develop 3D suspension culture methods allowing more reliable, simple, cost-effective and industrial-scale production of hPSC versus 2D systems. Free-floating 3D cultures (e.g. suspension culture in bioreactors) are expected to meet the high demand for cells needed for such applications. However, this approach can invoke problems with cell aggregation. Cells in the centre of these aggregates are generally underexposed to the medium, resulting in variable growth rates, apoptosis, uncontrolled differentiation and an eventual increase in heterogeneity^6^. Others have also explored the use of mechanical agitation (limited by shearing force), microcarriers, micropatterning and thermoreversible hydrogels^7, 8^. Despite their contribution to the field, each has its own limitations and struggle either to support rapid expansion or to prevent the formation of heterogeneous cell aggregates, leading to loss of pluripotency at higher cell densities^2, 8^. Chemical approaches to disrupt cell-cell contact in hPSC while maintaining pluripotency are an innovative approach to address this issue. A similar strategy has been previously described for mouse embryonic stem cells (mESC)^9, 10^. Here, we describe the novel use of a small molecule that reduces cell-cell adhesion, allowing the formation of smaller and more homogeneous hPSC aggregates while maintaining both pluripotency and viability, thereby improving scalability.

## Results

### (-)-ILV and OXS8360 induce iPSC spreading in 2D cultures

Disruption of cell-cell junctions is a requirement for epithelial cells to scatter^9^. With this in mind, a phenotypic screening assay was developed to identify small molecules causing hPSC colony disruption on the OX1-18 cell line^11^ in the first instance (Table S1). This identified the natural compound (-)-ILV (Fig. 1a), which allowed cells to separate from their neighbours in a concentration-dependent manner, with a minimal effective concentration of 1 µM (Fig. 1b). However, as (-)-ILV exhibited a reduction in cell viability over time (Fig. S1), an analysis of structural analogues of (-)-ILV was undertaken. One analogue, OXS8360 (Fig. 1a), induced iPSC cell scattering at 1 µM while maintaining high viability, as assessed by acridine orange. iPSC were cultured in the presence of (-)-ILV or OXS8360 and imaged after 3, 12 and 24h of treatment (Fig. 1b). Morphological changes were observable within 3h post treatment, with particularly pronounced separation of cells on colony peripheries, indicating the rapid effect of OXS8360. Colony disruption was more evident at 12h and reached a completely non-colony-based phenotype by 24h. Live imaging of OX8360-treated iPSC also showed a dynamically spreading phenotypic effect over 5 hours versus untreated cells (Video, S.I). Additionally, three genetically distinct healthy control donor iPSC lines (SFC840-03-06, SFC85403-02, and SFC856-03-04^12^) (Table S1) treated with either (-)-ILV or OXS8360 showed the same phenotypic effect, demonstrating the robustness of the effect (Fig. S2). Finally, the karyotypes of all four iPS cell lines remained stable over 10 passages of OXS8360-treatment (Fig. S3).

**Figure 1.**
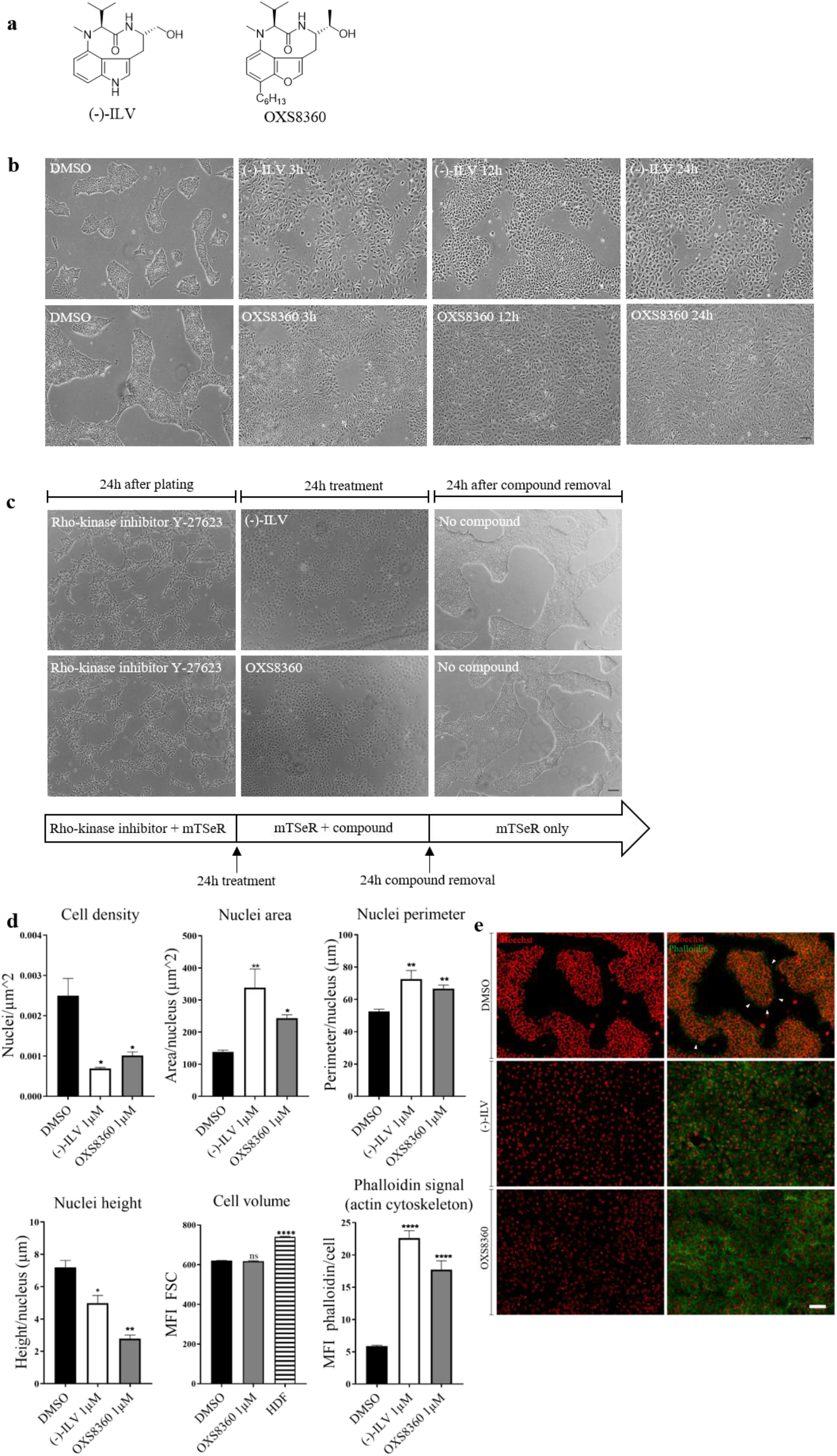
Phenotypic changes in iPSC induced by (-)-ILV and OXS8360 at 1μM. (**a**) Chemical structure of (-)-ILV and OXS8360. (**b**) Phase contrast images of (-)-ILV or OXS8360-treated iPSC at 3h, 12h and 24h. Colony disruption is observed 3h *post* treatment onwards. (**c**) Reversibility of the phenotypic effect induced by (-)-ILV and OXS8360. iPSC were plated with 10µM Rho-kinase inhibitor Y-27623 (upper panel) then treated for a subsequent 24h with (-)-ILV (left, middle panel) and OXS8360 (right, middle panel), then compounds were washed off (lower panel). (**d**) Effects of (-)-ILV and OXS8360 on cell density, nuclei area, perimeter, height, geometric mean forward scatter and actin cytoskeleton signal intensity. To determine nuclei area and perimeter, cells were fixed then stained with Hoechst for confocal microscopy analysis. To determine the nuclei height, z-stacks were imaged. Three images of different locations per condition were analysed with CellProfiler software. Statistical analysis was done using a One-way Anova with Dunett’s multiple comparison test. Error bars represent SEM. *P<0.05, **P<0.01, ****P<0.0001. (**d**) Immunofluorescence staining for actin filaments (green) and nuclei (Hoechst, red), showing cytoskeleton remodelling upon treatment with (-)-ILV or OXS8360. Scale bars, 100μm.

Next, the reversibility of (-)-ILV and OXS8360-induced morphological changes was assessed (experimental timeline schematically presented in Fig. 1c). 24h after plating (in medium containing Rho-kinase inhibitor Y27632 to prevent apoptosis during single cell passaging), iPSC started to form early colonies. Removal of Rho-kinase inhibitor and subsequent treatment with 1μM (-)-ILV or OXS8360 showed colony disruption 24h later. Finally, colony-based morphology was restored 24h after a compound-free medium change (Fig. 1c).

As (-)-ILV or OXS8360 treatment induced iPSC scattering, the resultant changes in cell morphology were quantified. Treatment with (-)-ILV or OXS8360 induced a significant reduction in cell density (nuclei/area) versus DMSO (3.5-fold and 2.7-fold, respectively). This was accompanied by a flattening of the phenotype, resulting in nuclei that were significantly greater in area and perimeter but significantly reduced in height (Fig. 1d). Geometric mean forward scatter (which reflects cell volume, obtained from flow cytometry datasets) showed no significant difference between OXS8360-treated and untreated iPSC (by comparison, human dermal fibroblasts (HDF) were significantly higher, Fig.1d), showing that overall cell volume was not changed, just a transition to a more flattened phenotype. Staining of iPSC actin filaments with phalloidin-iFluor 488 revealed cytoskeleton remodelling upon (-)-ILV or OXS8360 treatment. Control iPSC were characterized by few but organized actin filaments at cell-cell contacts, whereas (-)-ILV or OXS8360-treated iPSC had a significant increase (5.8 and 3.5 fold, respectively) in phalloidin signal, with F-actin arranged as stress fibres, indicative of weak cell-cell junctions^13, 14^ (Fig. 1d and 1e). These results show that both (-)-ILV and OXS8360 induce colony disruption involving substantial actin rearrangement in 2D-cultured iPSC, and that this phenotype is reversible.

### iPSC proliferate normally with OXS8360

As we had noted that (-)-ILV was toxic to iPSC, we explored the mechanism of cell death, and determined the concentration that inhibits cell survival by 50% for each compound. Activity assays for caspase 3 and 7 in whole cell lysates from iPSC treated over 24 h with concentrations ranging from 0.1 to 100 µM showed that (-)-ILV induced caspase 3 and 7 activation in a dose-dependent manner, with half maximal activity at 0.33 µM (Fig. S4). Half maximal activity for OXS8360-treated cells was at 89.1 µM, indicating much lower caspase induction. Flow cytometry for Annexin V-FITC and Propidium Iodide of (-)-ILV-treated cells showed that over 72h they shifted progressively from a population of healthy cells (Q4) to early apoptotic (Q3) and late apoptotic populations (Q2) (Fig. S5). These findings show that (-)-ILV exhibits high toxicity towards iPSC, therefore (-)-ILV was not used further, and only OXS8360 was taken forward for further studies.

To test whether iPSC proliferative ability would be affected following OXS8360 treatment, we examined expression of Ki-67, a protein that is present during all active phases of the cell cycle (G(1), S, G(2), and mitosis), but is absent from resting cells (G(0))^15^. Confocal imaging of untreated and OXS8360-treated iPSC for three passages revealed high levels of Ki-67 staining in both conditions (with HDF serving as a low-proliferating control) (Fig. 2a). Flow cytometry quantification of Ki-67-positive cells showed no significant difference between OXS8360 treated iPSC (99.4%±0.05; n=3), versus controls (99.0%±0.12; n=3), with HDF (12.3%±0.03, n=3) significantly lower than iPSC (Fig. 2b, 2c).

**Figure 2.**
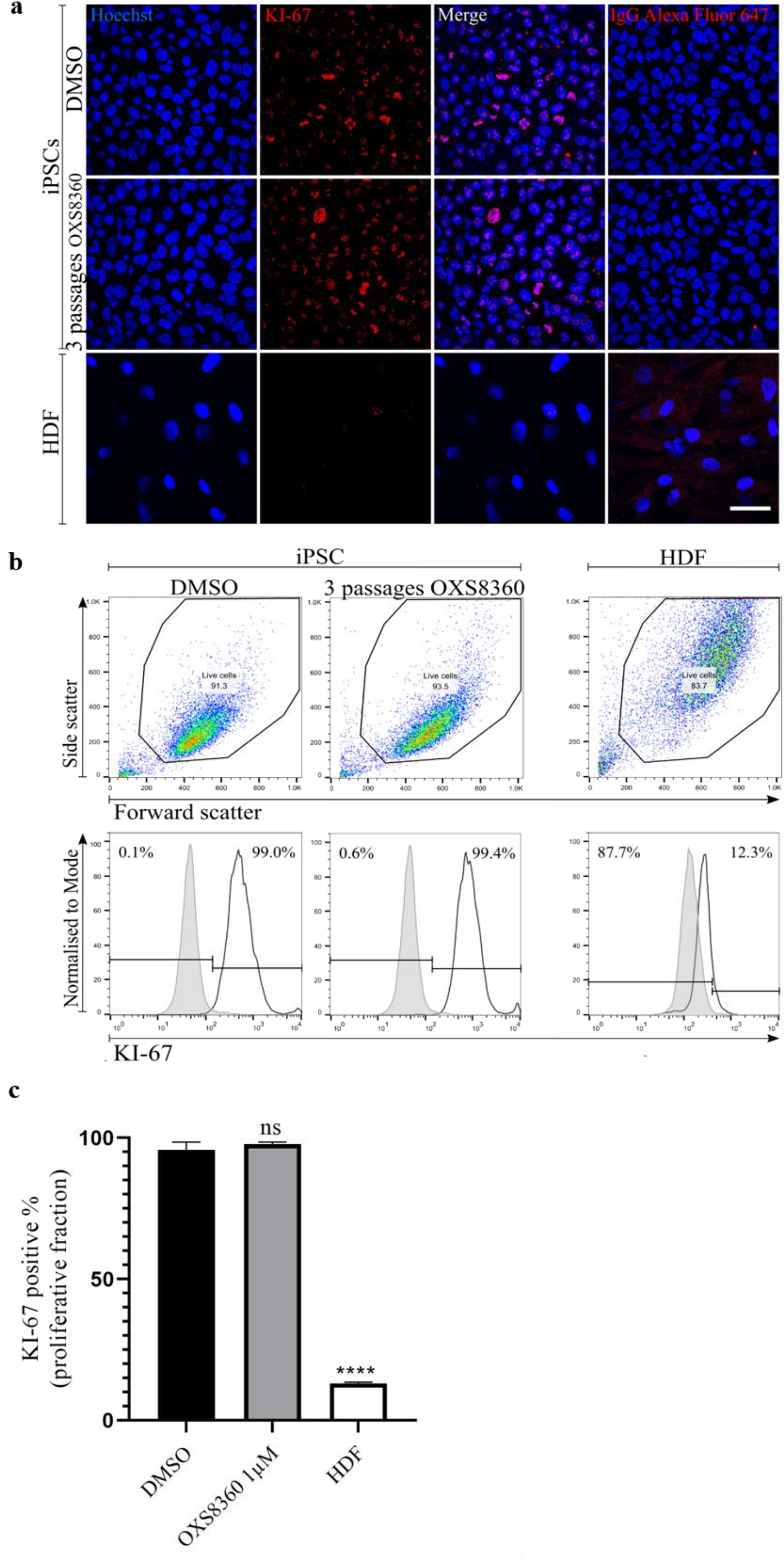
Assessing the proliferation capacity of OXS8360-treated iPSC through the expression of KI-67. (**a**) Immunofluorescence staining for KI-67 (red) shows a similar fraction of actively cycling cells in untreated versus iPSC treated with OXS8360 for three passages. Human dermal fibroblasts (HDF) serve as a slowly-proliferating comparison, with low KI-67 staining. Right hand panel, secondary antibody-only staining control. Scale bars, 50 µm. (**b**) Representative flow cytometry plots of KI-67^+^ cells. Black line and grey filled plots represent KI-67 stained population and negative control respectively. Debris and dead cells are gated out. Positive gate is set where negative control ≤1%. (**c**) Histograms for KI-67 positive populations in OXS8360-treated iPSC (mean±SEM: 99.4±0.05; n=3), DMSO-treated iPSC (mean±SEM: 99.0±0.12; n=3) and HDF (mean±SEM: 12.3±0.03; n=3).

### OXS8360-treated iPSC maintain pluripotent signatures

We next examined whether OXS8360 had any effect on the pluripotency signatures of iPSC treated over multiple passages. Firstly, untreated cells or cells treated for 3 passages were similarly Nanog^+^/Tra-1-60^+^/SSEA-3^+^/SSEA-4^+^ by flow cytometry (Fig. 3a), and negative for the mesenchymal surface marker CD44^16^. Furthermore, immunocytochemical analysis of cells confirmed Tra-1-60 and Nanog expression (Fig. 3b). iPSC treated with OXS8360 over 10 passages continued to express Tra-1-60 and Nanog, more consistently than the untreated controls, in all four genetically distinct cell lines (Fig. S6). Therefore, the expression of pluripotency markers in iPSC cultured with 1µM OXS8360 is stable over at least 10 passages.

**Figure 3.**
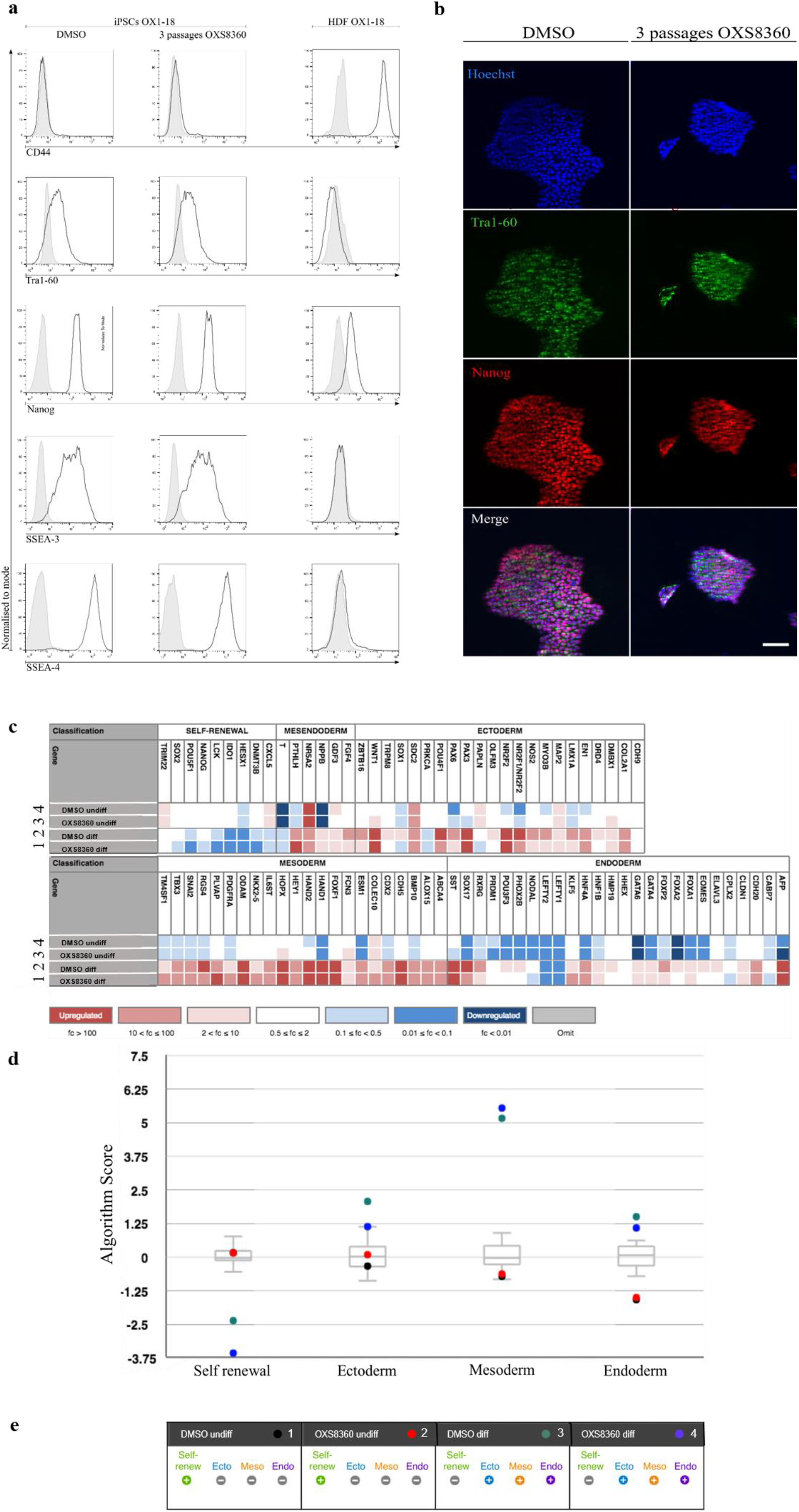
Assessing pluripotency signatures of OXS8360 treated iPSC. (**a**) Phenotypes of iPSC treated with DMSO or OXS8360, and HDF. Expression of the pluripotency markers Nanog, Tra-1-60, SSEA-3 and SSEA-4 and the fibroblast marker CD44 were measured by flow cytometry. The same gate to select for live cells was applied to all conditions. Histograms show antibody stained cells (black line plot) and relevant isotype control-stained cells (solid grey plot). (**b**) Immunocytochemistry staining showing expression of pluripotency markers Nanog and Tra-1-60 in untreated and OXS8360-treated iPSC. iPSC were treated over 3 passages with OXS8360 or DMSO followed by a 3 day culture without any compound prior to staining. Scale bar, 100μm. (**c**) Heat map groups genes according to signatures for pluripotency and trilineage differentiation for the 4 samples: (1) DMSO-treated undifferentiated, (2) OXS8360 treated for 3 passages undifferentiated, (3) DMSO treated then differentiated for 14 days, (4) OXS8360 treated then differentiated for 14 days. Each sample is normalised to a reference set of undifferentiated samples provided via the online software of the ScoreCard™ kit. Colour-coding indicates whether a gene is upregulated (red, ≥2), downregulated (blue, <0.5) or expressed at the same level (white, ≥0.5 and <2). (**d**) Box-and-whiskers plots indicate the reference signatures of undifferentiated samples as provided by the ScoreCard kit. 4 algorithm scores determine whether a sample is overall negative or positive for an entire differentiation or pluripotency signature. Algorithm scores ≥2 indicate upregulation, <0.5 downregulation and scores ≥0.5 and <2 are highly similar to a reference set of undifferentiated pluripotent stem cells. (**e**) Dots represent the algorithm scores of the included samples for each signature: DMSO-treated undifferentiated (black), OXS8360-treated undifferentiated (red), DMSO-treated and differentiated for 14 days (green), OXS8360-treated and differentiated for 14 days (blue).

Gene expression signatures can effectively report on the differentiation potential of hPSCs^17^, and ScoreCard, a qRT-PCR-based assay that evaluates pluripotency and differentiation signatures, offers a valuable quantitative approach for line-to-line comparison. Moreover, ScoreCard has been shown to be sensitive enough to compare the functional pluripotency of samples in distinct culture conditions, making it an appropriate genetic tool for our purpose^18^. Hence, Scorecard was used to test iPSC treated for three passages with OXS8360, to assess self-renewal signatures and undirected differentiation to the three germline lineages via embryoid bodies (EBs)^19, 20^ (imaged in Fig. S7). Both control DMSO and OXS8360-treated iPSC displayed expected expression signatures representing pluripotency, ectodermal, mesodermal and endodermal differentiation (Fig. 3c), expressed in an algorithm score against a reference dataset (Fig.3d), with consistent positive scores for pluripotency and negative scores for the three differentiation signatures as undifferentiated cells, and upregulated gene signatures for all three germ layers upon spontaneous differentiation (Fig.3e, Ct values Table S2).

### OXS8360-treated iPSC exhibit normal directed differentiation

We then assessed the capacity of iPSC treated with OXS8360 for three-passages to differentiate along defined pathways. iPSC differentiated to mesoderm-derived, primitive macrophages (pMac)^11, 21^ displayed expected morphology, including voluminous cytoplasm, membrane ruffles, and a proportion with an elongated spindle-like form^11^ (Fig.4a), together with characteristic macrophage cluster of differentiation (CD) markers CD11b, CD14 and CD45 (Fig. 4b). Directed differentiation of iPSC treated for three passages with OXS8360 to a neuroectodermal lineage (cortical neuron progenitors)^22^, gave rise to neuronal rosettes on day 16, followed by neurons visible at day 25 (Fig. 4c), which expressed cortical neuron markers TUJ1 and PAX-6 (Fig. 4d). Together, these results show that OXS8360-treated iPSC are competent at directed differentiation to key mesodermal and neurectodermal lineages.

**Figure 4.**
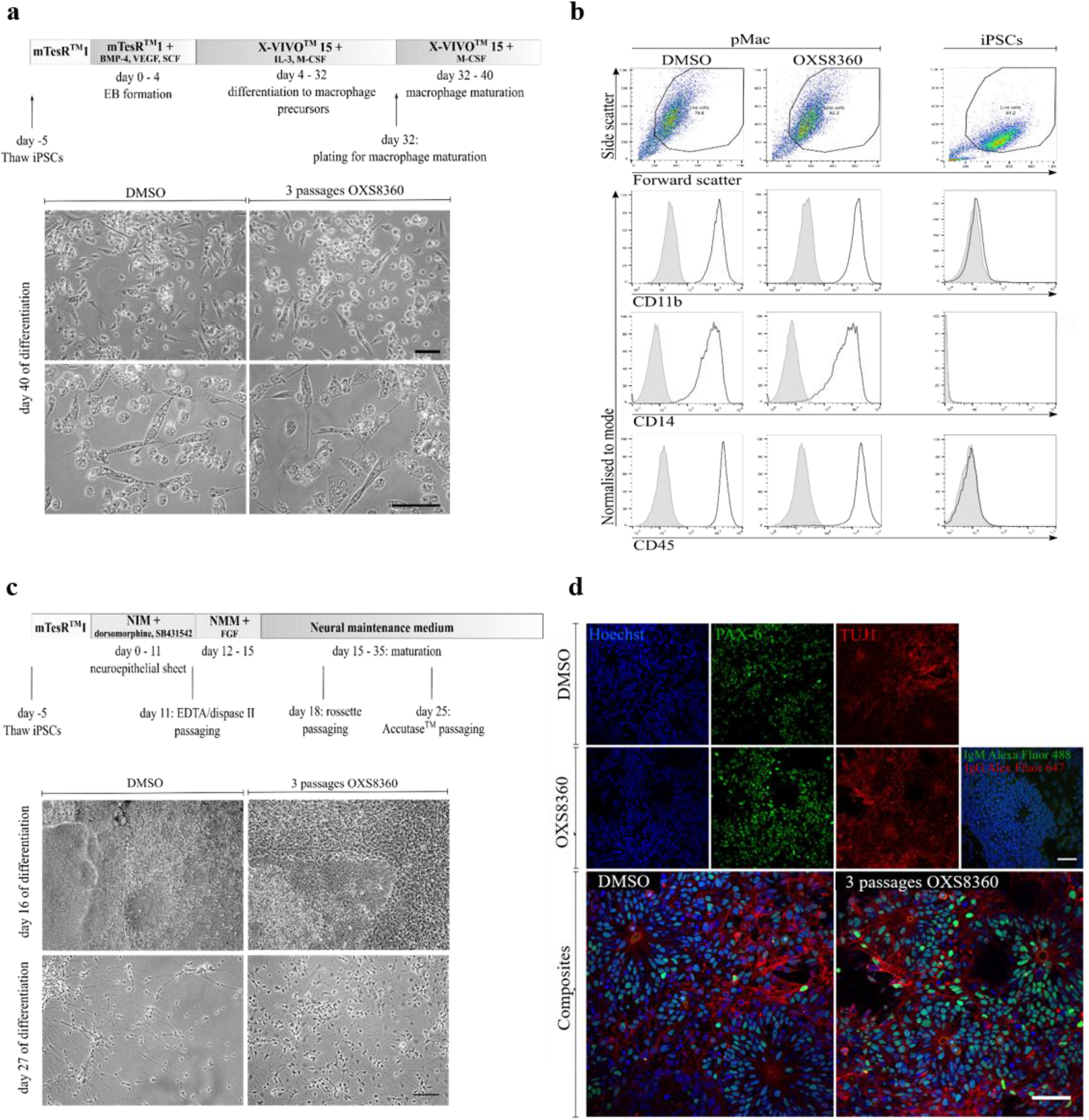
Assessing functional pluripotency through directed differentiation. (**a**) Morphological phenotypes of macrophages derived from DMSO and OXS8360-treated iPSC analysed using phase contrast microscopy. Shown are adherent macrophages (pMac) on day 40 of differentiation along the myeloid pathway. Scale bars, 100μm. (**b**) Flow cytometry showing the expression of CD11b, CD14 (both macrophage markers) and CD45 (pan-hematopoietic marker) at day 40 of differentiation. The same gate to select for live cells was applied to each condition. Histograms in each plot represent a population stained with the conjugated antibodies (black line plot) compared to its relevant isotype control (grey filled plot). (**c**) Differentiation of OXS8360 treated iPSC into cortical neurons. Phase contrast images of DMSO and OXS8360-treated iPSC on days 16 and 27. Scale bars, 50μm. (**d**) Immunocytochemistry staining for early neural markers PAX-6 and TUJ1 in cortical neuron cultures derived from DMSO and OXS8360-treated iPSC on day 25. Scale bars = 50μm.

### 3D culture of iPSC with OXS8360 enables formation of smaller aggregates

To assess whether the scattering of OXS8360-treated iPSC in 2D culture would lead to improved 3D culture, iPSC were seeded in six-well low attachment plates at 0.5×10^6^ cells per well (adapted from Zweigert e*t al.*^5^) and cultured for 10 passages in suspension, subculturing every 4 days with or without OXS8360. 4 days after the 10^th^ passage, untreated cells remained in large aggregates, whereas OXS8360 treated iPSC formed much smaller aggregates (Fig. 5a, Fig. S8). After 10 passages, Tra 1-60 and Nanog showed bimodal expression in the untreated cells, indicating a population of non-pluripotent cells, whereas OXS8360-treated cells retained a monomodal expression (Fig. 5b). Mean aggregate area was significantly smaller for OXS8360-treated (p < 0.001) versus untreated cultures (Fig. 5c). A three to six-fold expansion in seeded cells was observed during each passage for both conditions (Fig. S9). OXS8360-treated aggregates were much more easily dissociated to single cells as evidenced by significantly fewer residual cell clusters (14.5%±1.15, n=3, p<0.001) versus control cultures (33%±1.15, n=3) (Fig. 5d). Finally, the karyotype of iPSC treated with OXS8360 for 10 passages remained stable (Fig S3). These results show that iPSC can be 3D-cultured for extended periods of time in OXS8360 containing medium, reducing aggregation, maintaining pluripotency and facilitating dissociation.

**Figure 5.**
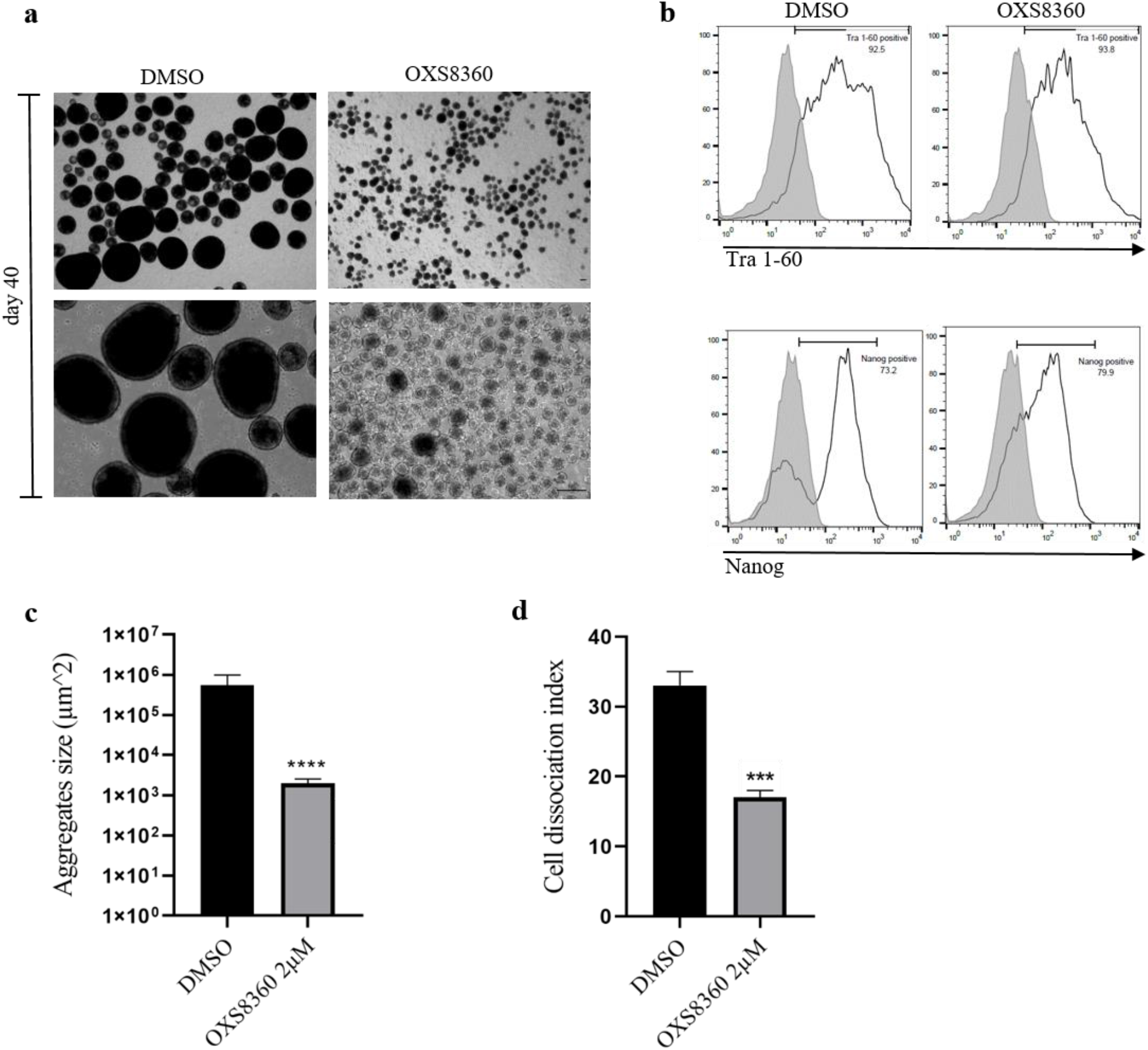
3D static suspension culture of iPSC. Cell-aggregation assay. Single iPSC were cultured for 10 passages with or without OXS8360 in static suspension culture in mTeSR and Rho-kinase inhibitor Y-27623 at a seeding density on each 4^th^ day passage of 1×10^6^ cells/mL. (**a**) Cell morphology on day 4 after the 10^th^ passage (phase contrast) shows smaller aggregates in presence of OXS8360 versus DMSO. Scale bar, 100µm. (**b**) Flow cytometry for Nanog and Tra1-60 in untreated and OXS8360-treated cells at passage 10. (**c**) Mean cross sectional area of the aggregates was determined with FIJI ImageJ. Note the formation of more homogenous aggregates in the presence of OXS8360 versus DMSO as indicated by error bars. (**d**) Cell dissociation assay. Aggregates were centrifuged, TryplE-dissociated and resuspended in culture medium. Cell dissociation index is expressed as the percentage of particles (cell clusters⩾4 cells) in total number of cells per well. Note the presence of fewer cell clusters following iPSC culture in the presence of OXS8360. Values are mean±SEM from three independent experiments. Statistical analysis was done using an unpaired t-test, ***P<0.001, ****P<0.0001.

### OXS8360 suppresses homophilic interactions between the extracellular domains of E-cadherin in iPSC

To explore the effect of OXS8360 on iPSC architecture, specifically E-cadherin-associated adherens junctions, OXS8360-treated iPSC were fixed and stained with anti E-cadherin mAb clones 36 and SHE78-7, commercially available antibodies that recognise an intracellular portion and the first extracellular domain (EC1) of E-cadherin respectively^23, 24^. Fluorescence intensity remained similar between treated and control cells when stained for intracellular E-cadherin (Fig. 6a). In contrast, and consistent with the observed disruption of iPSC colonies, the fluorescence intensity at cell-cell contacts revealed using the EC1 antibody recognising the extracellular domain was much weaker in OXS8360-treated cells than control cells.

**Figure 6.**
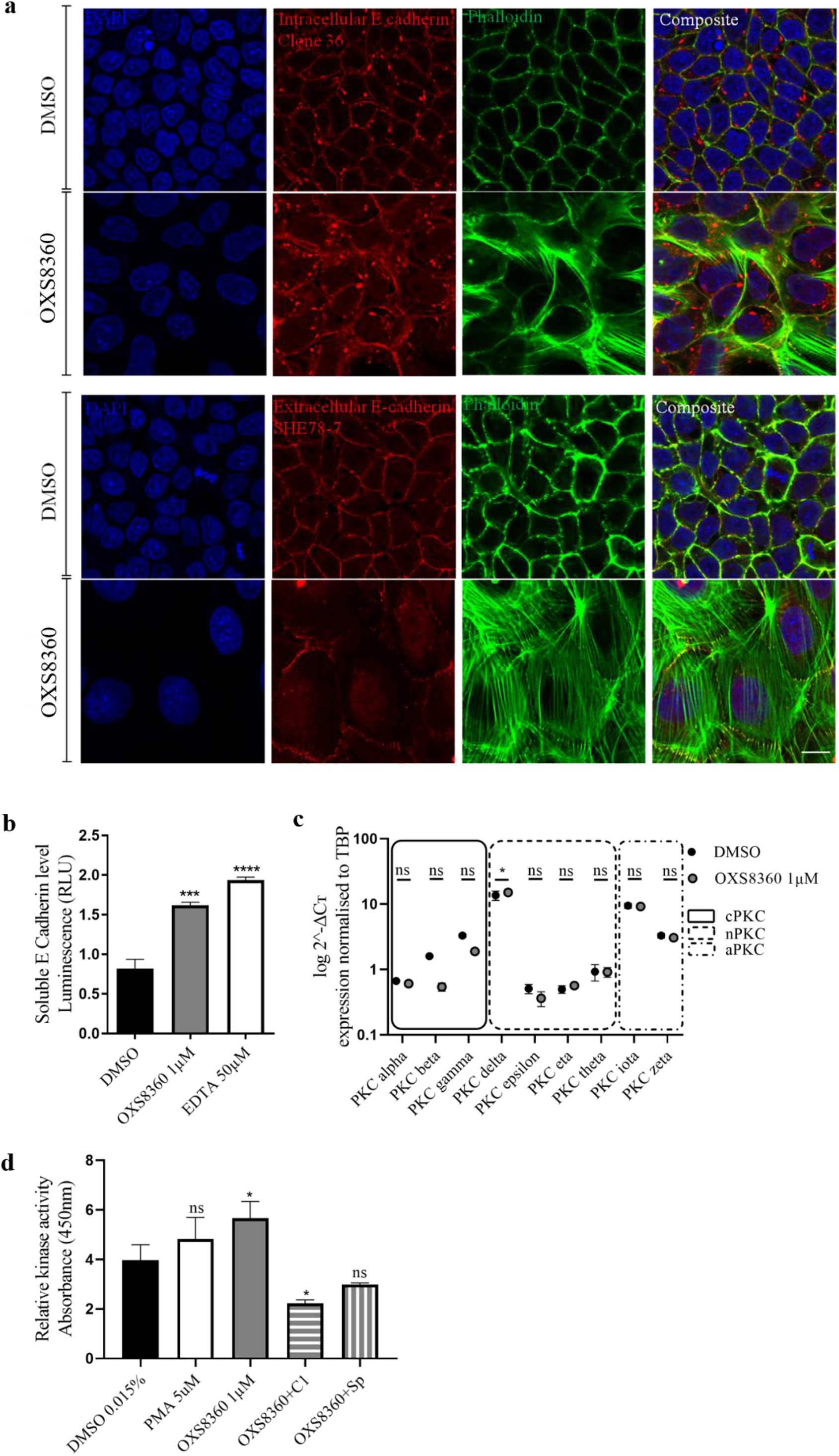
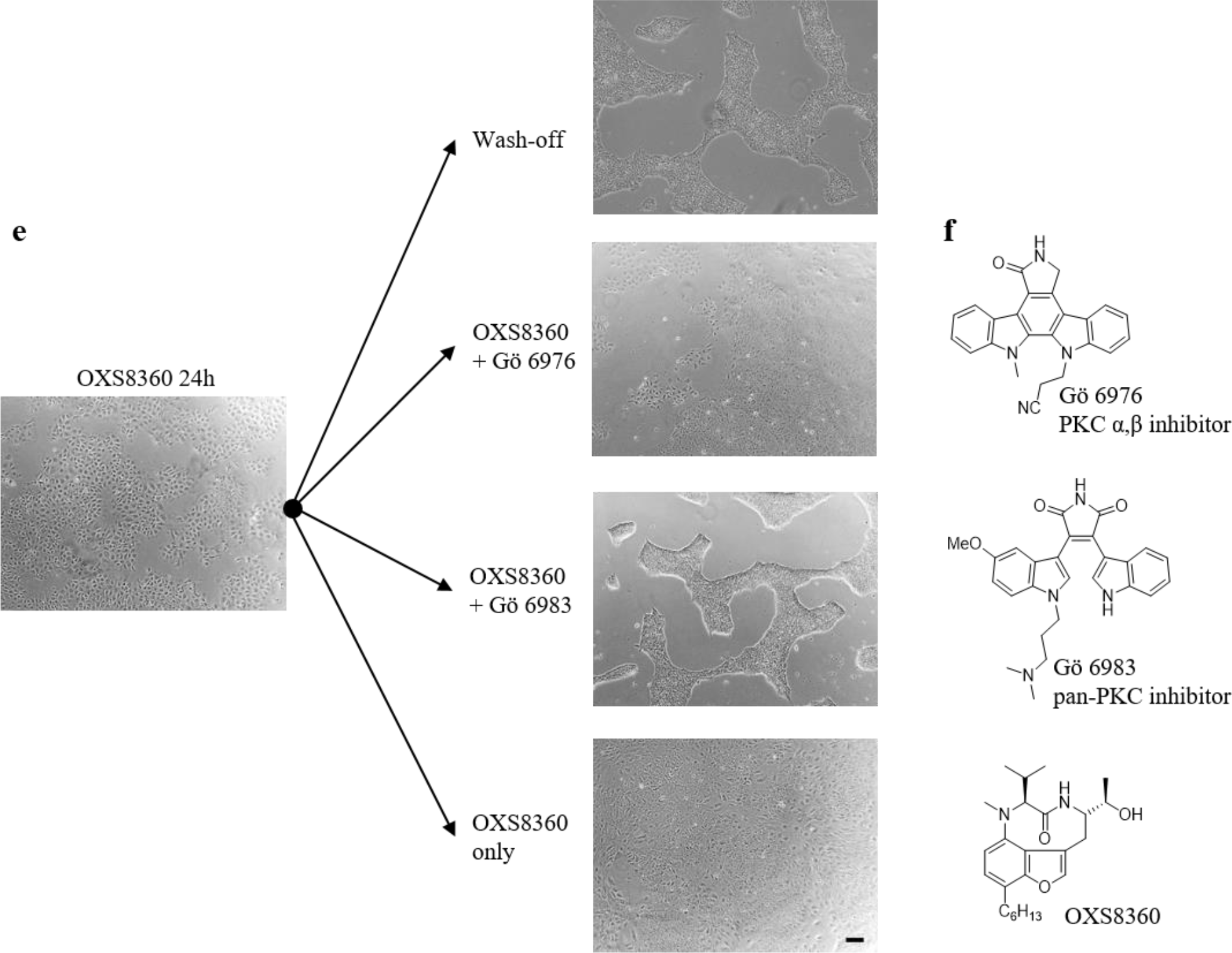
Mechanism of action of OXS8360. (**a**) Immunofluorescence staining for actin filaments (green), intracellular (antibody clone 36, red) and extracellular E-cadherin (antibody SHE78-7, red) in DMSO and OXS8360-treated iPSC. Nuclei were counterstained with Hoescht (blue). Cells were seeded overnight on µ-slides before being treated for 24h with or without OXS8360. Scale bar, 100µm. (**b**) Release of soluble E-Cadherin from cultured hiPSCs under OXS8360 and EDTA conditions. Monolayers of iPSC were cultured 24h prior to treatment. Statistical analysis was done using a one-way ANOVA with Dunnett’s multiple comparison test, ***P<0.001, ****P<0.0001, n=3. (**c**) PKC isoform expression in iPSC line OX1-18 upon treatment with OXS8360. qRT-PCR was performed to determine the relative expression levels of PKC isoforms normalised to TBP. iPSC were treated for 72h with OXS8360. All PKC isoforms are expressed in both treated and untreated samples. A significant difference could only be observed for the expression of PKC δ. Statistical analysis was done using a one-way ANOVA with Sidak’s multiple comparison test, ns P>0.05, *P<0.05, n=3. (**d**) Active PKC protein levels in iPSC following treatment with PMA, OXS8360 and/or PKC inhibitors (C1 and Sphingosine (Sp)). Cells from PMA or OXS8360 treatment group alone were homogenised prior to co-treatment with PKC inhibitors. Statistical analysis was done using a one-way ANOVA with Dunnett’s multiple comparison test, ns P>0.05, *P<0.05, n=3. (**e**) Phase contrast images of iPSC treated with 1 µM OXS8360. After 24h, the medium was changed and the treatment was continued either in a combinatorial approach with the PKC inhibitors Gö6976 and Gö6983 (both at 1 µM) either with OXS8360 (positive control) or medium alone (negative control). iPSC treated with OXS8360 and the PKC inhibitor Gö6983 displayed colony-based morphology. Co-treatment with Gö6976 did not abrogate the colony-spread phenotype. Scale bars, 100 µm. (**f**) Chemical structures of PKC inhibitors Gö6976 (α, β) and Gö6983 (pan-PKC) and OXS8360.

Separation of adjacent epithelial cells has been shown to be induced by disruption of cell-cell E-cadherin interactions. Several examples of mechanisms resulting in loss of this protein’s surface expression include internalization of E-cadherin, proteosomal degradation, and proteolytic cleavage^25^. Cleavage of mature E-cadherin (120 kDa), results in the shedding of a soluble 80 kDa E-cadherin fragment. Therefore, we sought to measure its presence in the supernatant of both treated and untreated iPSC. Cells were plated and grown for 24 hours prior to treatment with 0.1% DMSO (negative control), 50 µM ethylenediaminetetraacetic acid (EDTA) (used as a positive control shown to enhance E-cadherin cleavage^26^) or 1 µM OXS8360 for 24 hours. Soluble E-cadherin released from cultured cells significantly increased in the presence of OXS8360 (Fig. 6b), as measured by ELISA, consistent with the adhesion-inhibitory effects.

### PKC modulates iPSC spreading induced by OXS8360

As OXS8360 has been reported to be a protein kinase C (PKC) activator, binding to the conventional (α, β, γ) and novel (δ, ε) PKC isoforms^27^, we hypothesised that the observed iPSC phenotype may be mediated through this pathway. A comparison with structurally unrelated PKC activators was first carried out. Treatment of iPSC with commercially available phorbol esters (known to be PKC activators^28^) gave a similar effect on colony disruption to OXS8360 (Fig. S10). We next determined the expression levels of PKC isoforms in iPSC, which have been reported to exhibit tissue-specific levels of expression^28^ All conventional, novel and atypical PKC isoforms (cPKC, nPKC, aPKC) were expressed in the iPSC line OX1-18, with PKC δ, ι and ζ the most highly expressed. With the exception of PKCδ, no significant differences in the expression of isoforms were observed upon 72h treatment with OXS8360 compared with untreated cells (Fig. 6c).

In addition, using a relative kinase assay, we observed that 1 µM OXS8360 induced a statistically significant increase in PKC enzymatic activity in iPSC (Fig. 6d).

Finally, two pharmacological PKC inhibitors, namely Gö6976 and Gö6983, were tested for their ability to block OXS8360-induced morphological change in iPSC. These inhibitors have been shown to inhibit PKC isoforms at nanomolar concentrations^29, 30^. Gö6976 has been reported to be a specific inhibitor of the conventional PKC isoforms α and β whereas Gö6983 acts as pan-PKC inhibitor with more affinity for conventional and novel isoforms than atypical isoforms^29, 30^. While blocking the PKC pathway with conventional PKC inhibitor Gö6976 had no effect, pan-PKC inhibitor Gö6983 reversed the effect of OXS8360 on colony disruption (Fig. 6e), suggesting the effects of OXS8360 may be mediated via unconventional PKC isoforms.

Taken together this data suggests that the effect of OXS8360 may be mediated at least in part through the activation of novel PKC isoforms.

## Discussion

This study has identified a small molecule, OXS8360, which overcomes some of the limitations in iPSC culture by reversibly disrupting 2D colonies and 3D aggregates, thereby allowing more homogeneous cultures. Our phenotypic studies were reproducible across four genetically distinct iPS cell lines, overcoming inherent iPSC line-to-line variability and molecular heterogeneity^31^.

OXS8360-treated 2D-cultured cells displayed remodelling of the actin cytoskeleton and less tightly-packed colonies. However, note that we could not confirm total contact-independence. OXS8360-treated 3D-cultured cells displayed smaller, looser aggregates, resulting in a more homogeneous cell population, with improved dissociation capacity. This overcomes several limitations of current 3D aggregate systems, especially transport of nutrients and small molecules that play an important role in pluripotency maintenance^6, 32^.

OXS8360-treated iPSC maintained a highly proliferative self-renewal capacity^33^ (assessed by Ki-67 staining and cumulative cell-count), and a pluripotent signature, shown by expression of widely accepted markers^34^, ScoreCard assay, undirected differentiation into all three germ layers, and directed differentiation to mesodermal (macrophages) and neurectodermal (cerebral cortical neurons) lineages. Extended culture for 10 passages demonstrated that OXS8360-treated iPSC preserved their key iPSC characteristics, including karyotypic stability. Therefore, OXS8360 is fully compatible with iPSC functionality over extended passaging.

To elucidate the mechanism of action of OXS8360, and consistent with previous reports^27^, we showed that OXS8360 increased PKC activity in iPSC. PKC is a pleiotropic enzyme, but also mediates the tumour-promoting activity of phorbol esters and certain teleocidin natural products, including (-)-ILV^35^. The tumor-promoting potential of the phorbol ester TPA is context dependent^35^, and structure-activity-relationship analysis of (-)- ILV has demonstrated that tumour-promoting activity is not observed across all related structures and not necessarily dependent on PKC activation^36^. Furthermore activation of PKC by other chemotypes (e.g. bryostatin) has been demonstrated to lack tumour-promotor activity^37^. PKC activators have not been reported as tumour *initiators*, and importantly in our study we see no evidence of any effect on genome integrity or proliferation rate following OXS8360-treatment.

All cPKCs, nPKCs and aPKCs isozyme transcripts were detectable in iPSC. However, it remains unclear whether OXS8360 exerts its effects through specific PKC isozymes. Moreover, specific PKC isoforms^38^ or exposure to activators^35^ can have opposing effects in different cell types, depending on the stimulus and PKC intracellular localisation. Therefore we compare our results here with other reports studying PKC isoform effects in epithelial cell types. The effect of OXS8360 on iPSC was not abrogated by Gö6976 (a PKC α and β inhibitor), however, this does not rule out an effect mediated by PKCγ. In rabbit lens epithelial cells, PKCγ has been reported to play a role in the regulation of the gap junction protein, connexin 43^39^. Of the nPKCs, PKCε has been shown to suppress adherens junctions through adducin^40^, weaken tight junctions through claudin-4^41^ and promote epithelial-mesenchymal transition (EMT)^42^. PKCδ has been shown to modulate cell scattering in MDCK cells^23^ and human keratinocytes^43^. Both studies suggest that PKCδ might interfere with homophilic interactions between E-cadherin ectodomains, thus suppressing adherens junctions. Meanwhile, Oh *et al.*^44^ reported the involvement of PKCδ in the regulation of peripheral actin organization and cell-cell contacts in the epithelium. Interestingly, PKCδ was found to be the most highly expressed isoform in iPSC.

OXS8360-treated iPSC shed more E-cadherin into the medium, and the E-cadherin ectodomain (immunostaining with antibody SHE78-7) decreased, whereas the cytoplasmic domain of E-cadherin (antibody clone 36) was unchanged, as previously observed^23^. It is possible that PKC directly phosphorylates the cytoplasmic domain of E-cadherin causing a conformational change on the ectodomains, disfavoring the homophilic binding of the ectodomains of E-cadherin, identified as cleavage sites for proteases^45, 46^. Alternatively, other proteins involved in the assembly of adherens junctions may be phosphorylated by PKC activation indirectly. PKC-mediated phosphorylation of β-catenin negatively regulates the Wnt/ β-catenin pathway leading to disconnection of its interaction with cytoplasmic E-cadherin^47^. Overall we propose that activation of PKC leads to weakening of cell-cell junctions and shedding of soluble E-cadherin, although further studies will be necessary to clarify which PKC isoform or isoforms are involved and the intermediate mechanistic steps.

## Conclusion

In summary, we have identified a small molecule additive, OXS8360, which enables more uniform 2D and 3D culture of iPSC without compromising potency or genetic integrity. In 3D culture systems, our approach reduces the number of culture components needed, as no hydrogel or microcarrier-based culture is required to control aggregate size. The looser and smaller aggregates facilitate maintenance and expansion of iPSC while also overcoming issues of diffusion and homogeneity. This approach will improve the efficiency and reproducibility of hPSC culture at scale.

## Statistical analysis

GraphPad Prism was used for statistical analysis. One-way ANOVA with Sidak’s or Dunnett’s multiple comparisons or paired two-tailed t test were used as indicated. Values are indicated in figures as ^∗^p < 0.05, ^∗∗^p < 0.01, ^∗∗∗^p < 0.001, ^∗∗∗∗^p < 0.0001, and n.s. (not significant).

## Methods

### Cell lines and culture

This study was carried out using four reprogrammed iPSC lines (Table S1), with human dermal fibroblasts (HDF) as a control where necessary. The iPSC and HDF were all derived in Oxford from healthy control donors recruited through the Oxford Parkinson’s Disease Centre - participants were recruited having given signed informed consent, which included derivation of hiPSC lines from skin biopsies (Ethics Committee, National Health Service Health Research Authority, NRES Committee South Central, Berkshire, UK, REC 10/H0505/71). All experiments were performed in accordance with UK guidelines and regulations and as set out in the REC and the iPSC lines used have all been published previously (Table S1).

iPSCs were maintained in 2D feeder-free conditions on hESC-qualified Matrigel (Corning), or Geltrex (Life Technologies) in complete mTeSR™1 (Stem Cell Technologies) supplemented with 100 units/mL of penicillin-streptomycin (Gibco™) at 37°C in a humidified atmosphere with 5% CO2. At approximatively 80% confluence, cells were dissociated for 5 min at 37°C in prewarmed TrypLE Express (Gibco™). TrypLE Express was then diluted 1:10 in prewarmed 1x Dulbecco’s phosphate-buffered saline (DPBS), and cells centrifuged for 5 min at 400 rcf. Supernatant was removed and cell pellets were gently resuspended to give a single cell suspension in medium supplemented with 10 μM ROCK inhibitor Y-27632 (Sigma-Aldrich). Daily medium change was performed by replacing the medium with fresh mTeSR™ without ROCK inhibitor Y-27632.

For static suspension culture^i^, 6-well plates with repellant surfaces were used (Greiner Bio-One™). 3 mL of a cell suspension at a density of 0.33×10^6^ cells/mL were supplemented with 100 units/mL penicillin-streptomycin and 10 µM ROCK inhibitor Y-27632. Aggregates were allowed to form at 37°C in a humidified atmosphere with 5% CO2 over 4 days with no medium change. On day 4 of suspension culture, aggregates were transferred to 15-mL tubes. Supernatant was removed after centrifugation at 400 rcf for 5 min. This was followed by incubation in 1 mL of TrypLE for 5 min in a water bath set at 37°C. Cells were centrifuged at 300 rcf for 4 min to remove the supernantant and resuspended in 1 mL of mTesR™ prior to filtration through a 30 μm sieve. The membrane was washed with 1 mL mTeSR™ and cells were resuspended in the appropriate volume of ROCK inhibitor Y-27632 supplemented mTeSR1™. A dedicated incubator was used to prevent inadvertent agitation of the cultures through door opening/closing.

When treated with compounds for experiments, cells were cultured in the presence of 0.1% DMSO, 1 µM (-)-ILV, or 1 µM OXS8360 unless stated otherwise. Cell phenotype was monitored by using phase contrast microscopy with an EVOS XL cell imaging system.

HDFs were cultured in T-25 flasks (Corning) at 37°C and 5% CO_2_ with 6 mL HDF medium consisting of 1x Advanced DMEM (Gibco™), 1x Glutamax™ (Gibco™), 100 μM β-mercaptoethanol (Gibco™), 0.1x fetal bovine serum (Sigma). Medium was changed every 4 days and cell passaging was as described for iPSCs.

### Caspase activity assay

Measurements of caspase activities in cells were performed using the commercially available Caspase-Glo 3/7 Assay (Promega, Madison, WI) according to the manufacturer’s instructions. A previous experiment for the determination of the cell density revealed an optimal linear range of the assay at 10.000 cells/well in a 96 well format.

### Annexin V assay

iPSCs were seeded at a density of 1×10^5^ cells/well onto a 24-well plate, and treated for 24h, 48h and 72h with 1 µM of (-)-ILV or 0.1% DMSO and apoptotic cells were analyzed by using Annexin V FITC apoptosis detection kit (Abcam). Following treatment, cells were washed with 500 µL Hank’s Balanced Salt Solution (HBSS, Gibco™) and dissociated as described above. Centrifugation was 400 rcf for 5 min and pellets resuspended in a 100 μL 1x binding buffer. Further incubation was performed with 5 μL of FITC-conjugated Annexin V and 5 μL of propidium iodide for 15 min in the dark. Prior to analysis, 200 μL of binding buffer was added to each sample. Quantification was realized by flow cytometry and data analyzed using FlowJo software.

### Flow cytometry

For assessing cell surface antigens, 30 min staining on ice was carried out on fresh cells in cold FACS buffer consisting of PBS, 10 μg/mL human IgG (Sigma), 1% FBS (Hyclone) and 0.01% sodium azide prior to fixation. For intracellular antigens, staining was carried out on cells fixed with 4% paraformaldehyde (PFA) in PBS for 10 min and stored in methanol at −20°C for up to 4 weeks. Cells were stained at a density of 1×10^6^ cells/mL in a total of 100 μL FACS buffer with the appropriate antibody or isotype-matched control in a V-bottom 96-well plate. For two-colour staining, two antibodies or two isotype controls (attached to different fluorophores) were added together. Following an incubation of 90 min at room temperature, cells were washed three times with FACS buffer. When using unconjugated antibodies, a further incubation of 40 min, at room temperature, was performed with a secondary antibody. Samples were washed another 3 times and analysed using FACS Calibur flow cytometer (Becton Dickinson). Data were analysed using FlowJo software and antibodies used are described in the table below.

| Product | Host | Isotype, conjugate<br>fluorophore | Supplier | [c] | Catalogue<br>number |
| --- | --- | --- | --- | --- | --- |
| Anti-CD44 | Mouse | IgG<sub>1</sub>, FITC | Life Technologies | 1:40 | 11044182 |
| Anti-SSEA3 | Rat | IgGM, Alexa Fluor 488 | Biolegend | 1:80 | 330306 |
| Anti-SSEA4 | Mouse | IgG<sub>3</sub>, APC | R&D | 1:20 | FAB1435A |
| Anti-Tra1-60 | Mouse | IgM, PE | Biolegend | 1:10 | 330609 |
| Anti-Tra1-60 | Mouse | IgM, Alexa Fluor 488 | Biolegend | 1:10 | 330613 |
| Anti-Nanog | Rabbit | IgG, Alexa Fluor 647 | Cell Signaling | 1:10 | 5448S |
| Anti-KI-67 | Mouse | IgG<sub>1</sub> | Merck | 1:400 | MAB4190 |
| Anti-CD11b | Mouse | IgG<sub>1</sub>, APC | Biolegend | 1:25 | 301309 |
| Anti-CD14 | Mouse | IgG<sub>1</sub>, PE | ImmunoTools | 1:25 | 21620144 |
| Anti-CD45 | Mouse | IgG<sub>1</sub>, APC | ImmunoTools | 1:25 | 21270456 |
| Isotype control | Mouse | IgG<sub>1</sub>, FITC | Biolegend | 1:50 | 400107 |
| Isotype control | Rat | IgM, Alexa Fluor 488 | Biolegend | 1:100 | 400811 |
| Isotype control | Mouse | IgG<sub>3</sub>, APC | R&D | 1:20 | IC007A |
| Isotype control | Mouse | IgM, Alexa Fluor 488 | Biolegend | 1:10 | 401617 |
| Isotype control | Rabbit | IgG, Alexa Fluor 647 | Cell Signaling | 1:10 | 2985S |
| Secondary antibody | Donkey | IgG, anti-mouse Alexa Fluor 647 | Invitrogen | 1:1000 | A-31571 |
| Secondary antibody | Goat | IgM, anti-mouse, Alexa Fluor 488 | Invitrogen | 1:1000 | A-21042 |
| Secondary antibody | Goat | IgG<sub>2a</sub>, anti-mouse, Alexa Fluor 647 | Invitrogen | 1:1000 | A-21241 |
| Secondary antibody | Goat | IgG, anti-rabbit, Alexa Fluor 647 | Invitrogen | 1:1000 | A-21244 |

### Undirected differentiation

iPSC were washed with PBS and harvested by incubating the cells for 5 min at 37°C with 1 mL warm TrypLE Express (Gibco™). The cells and TrypLE were well mixed into a single cell suspension by pipetting up and down and collected in a 15 mL centrifuge tube and diluted 1∶10 with PBS. Cells were counted and spun down. After centrifugation PBS was aspirated and the cell pellet was tapped loose and resuspended in mTeSR™-1 spin-EB medium consisting of mTeSR™-1 (Stem Cell Technologies), supplemented with 1 mM Rock-inhibitor (Y27632, Calbiochem). For AggreWells™800 (Stemcell Technologies, 300 micro-wells), plates were first prepared After preparing the plate, 1 mL of 4×10^6^ PSC were added per well. The plate containing PSC and 2 mL of spin-EB medium per well was centrifuged at 800 rpm for 3 minutes. The plate was examined under the microscope to verify cells were evenly distributed among the micro-wells. Very gently the plate was put into the incubator and left for four days. The EBs were fed daily with spin-EB medium (first brought to RT), by gently aspirating 1 mL medium using a p1000 Gilson and very gently adding 1 mL fresh spin-EB medium in a drop-wise manner down the side of the well so the EBs were not washed out of the microwells. This wash was repeated to achieve a 75% medium change overall. To harvest EBs at day 4, the contents of the wells were pipetted up and down several times using a 5 mL serological pipette to dislodge the EBs from the micro-wells. The contents were taken up and transferred onto a 40 µM cell strainer inverted over a 50 mL centrifuge tube. The inverted strainer, with the EBs balanced on top, was carefully inverted over onto a new 50 mL centrifuge tube so that the EBs were now at the bottom of the strainer and could be collected into the new 50 mL tube, by passing through 4 mL of relevant differentiation medium. The strainer was held at an angle to facilitate the collection of EBs. The EBs for each condition were then plated split into half and plated onto a Geltrex pre-coated 60mm petri dish (Corning) for the TaqMan hPSC ScorecardTM Panel assay.

### TaqMan hPSC Scorecard™ Panel assay

To form EBs, the manufaturer’s protocol described in the user guide for the TaqMan hPSC ScorecardTM Panel (Applied Biosystems™) was adapted. EBs were generated according to the EB-spin formation protocol and harvested on day 4 as outlined above. Between day 4 and 14, a 75% medium change was performed only every other day. On day 14, each cell culture dish was harvested. The culture medium was removed and cells were gently washed in 5mL PBS for 2 minutes. 700 μl RLT buffer (Qiagen) was pipetted onto each plate supplemented with 1% of β-mercaptoethanol (Sigma) to reduce RNase activity. Upon cell lysis, a cell scraper (Sarstedt) was used to collect the remaining cell content. After careful homogenisation by using a Gilson p1000, the total of 700μl slurry was transferred in aliquots of 350 μl onto two QIAGEN shredder membranes (Qiagen), and stored at −80°C.

### Directed differentiation to macrophages

iPSCs were differentiated to macrophages as previously described by van Wilgenburg *et al.*^ii^. Briefly, 3 × 10^6^ iPSCs were seeded into an Aggrewell 800 well (STEMCELL Technologies) to form EBs, in mTeSR1 and fed daily with medium plus 50 ng/mL BMP4 (Peprotech), 50 ng/mL VEGF (Peprotech), and 20 ng/mL SCF (Miltenyi Biotec). Four-day EBs were then differentiated in T175 flasks (150 EBs) in X-VIVO15 (Lonza), supplemented with 100 ng/mL M-CSF (Invitrogen), 25 ng/mL IL-3 (R&D), 2 mM Glutamax (Invitrogen), 100 U/mL penicillin and 100 μg/mL streptomycin (Invitrogen), and 0.055 mM β-mercaptoethanol (Invitrogen), with fresh medium added weekly. Macrophage precursors (pMacpre) emerging into the supernatant after approximately 1 month were collected weekly and differentiation cultures replenished with fresh medium. Harvested cells were strained (40 μm, Corning) and plated onto tissue-culture treated plastic at 100,000 per cm^2^ and differentiated for 8 days to adherent macrophages (pMac) in X-VIVO15 with 100 ng/mL M-CSF, 2 mM Glutamax, 100 units/mL penicillin-streptomycin.

### Directed differentiation to cortical neuron progenitors

iPSCs were differentiated to cortical neuron progenitors (NPCs)^iii^ with the following modifications: feeder-free iPSCs were plated onto Matrigel-coated 6-well plates, with neural induction for 12 days using dual SMAD inhibition; after replating the neuroepithelial sheet on precoated wells with 20 µg/mL of laminin at day 12 using dispase II, 20 ng/mL fibroblast growth factor 2 (FGF2, R&D) was added to neural maintenance medium (NMM) from days 12 to 15. Thereafter, a full medium change was performed every other day. On day 18, newly formed rosettes were passaged either with dispase II or manually. For the latter, rectangles containing rosettes were cut with a needle, and in case of large size, scored into multiple rectangles. The cell sheets were then lifted manually, collected with a Gilson p200 and transferred in 2mL of NMM onto a new well precoated with 20μg/mL laminin. Between day 18 and 25, a full medium change was performed every other day, and cells were stained on day 25 for confocal imaging.

### Immunocytochemistry

Undifferentiated iPSCs on Geltrex coated µ-slides were prepared for immunofluorescence microscopy as followed. E-cadherin expression was monitored for cells seeded overnight prior to 24h treatment with 0.1% DMSO, 1 µM ILV or 1 µM OXS8360. When looking at the expression of KI-67, Tra 1-60 or Nanog cells were treated for 3 passages. Treated iPSCs were fixed with 4% PFA for 10 minutes at room temperature, permeabilised and blocked with 10% goat serum and 0.3% Triton X-100 in PBS overnight at 4°C prior to washing 3 times with PBS and 0.3% Triton X-100. Anti-E-cadherin (2.5 µg/mL, clone 36, BD Transduction Laboratories or SHE78-7, Invitrogen), anti-KI-67 (1:400, Merck), anti-Tra-1-60 PE (conjugated, 1:10, Biolegend), anti-Nanog Alexa Fluor 647 (conjugated, 1:10, Biolegend) antibodies were diluted in antibody solution (PBS, 1% bovine serum albumin (BSA), 0.1% Triton X-100) and incubated overnight at 4°C. When using unconjugated antibodies, wells were washed prior to applying Alexa Fluor 647 (goat-anti-rabbit, IgG, 1:1000) diluted in antibody solution (PBS, 1% BSA, 0.3% Triton-X) for 2 hours at room temperature.

For cortical neuronal progenitors, pretreated cells with 0.1% DMSO or 1 µM OXS8360 were passaged on day 18 onto precoated µ-slides and grown until day 25. Cells were fixed with 4% PFA for 10 minutes at room temperature, permeabilised and blocked with PBS, 1% donkey serum/1% goat serum, 0.3% Triton-X overnight at 4°C. Anti-PAX-6 (1:300, Covance) and anti-TUJ-1 (1:200, Covance) were used as primary antibodies, and Alexa 488 (goat-anti-mouse, IgM, 1:1000) or Alexa Fluor 647 (goat-anti-rabbit, IgG, 1:1000) were stained for visualization.

After washes, nuclei were counterstained with Hoechst 33342 or DAPI for 30 minutes at room temperature. Fluorescent images were acquired using the FV1200 (Olympus) confocal microscope. All antibodies used are described in the table below.

| Product | Host | Isotype, conjugate fluorophore | Supplier | [c] | Catalogue number |
| --- | --- | --- | --- | --- | --- |
| Anti-E-cadherin, clone 36 | Mouse |  | BD Transduction Laboratories | 2.5 µg/mL | 610182 |
| Anti-E-cadherin, clone SHE78-7 | Mouse |  | Invitrogen | 2.5 µg/mL | 13-5700 |
| Anti-Tra1-60 | Mouse | IgM, PE | Biolegend | 1:10 | 330609 |
| Anti-Nanog | Rabbit | IgG, Alexa Fluor 647 | Cell Signalling | 1:10 | 5448S |
| Anti-KI-67 | Mouse | IgG<sub>1</sub> | Merck | 1:400 | MAB4190 |
| Anti-PAX-6 | Rabbit | IgG (polyclonal) | Covance | 1:300 | *PRB-278P* |
| Anti-TUJ1 | Mouse | IgG<sub>2a</sub> | Covance | 1:200 | MMS-435P |
| Secondary antibody | Donkey | IgG, anti-mouse Alexa Fluor 647 | Invitrogen | 1:1000 | A-31571 |
| Secondary antibody | Goat | IgM, anti-mouse, Alexa Fluor 488 | Invitrogen | 1:1000 | A-21042 |
| Secondary antibody | Goat | IgG, anti-rabbit, Alexa Fluor 647 | Invitrogen | 1:1000 | A-21244 |

### Enzyme-linked Immunosorbent Assay for soluble E-Cadherin

Cells were seeded in a 6-well plate at a density of 1 x 10^6^ cells per well for 24 h prior to starting any treatment. Medium was removed and a further incubation was realized in presence of 0.1% DMSO, 1 µM OXS8360 or 50 µM EDTA in fresh mTeSR™. Medium from each condition was removed and E-cadherin concentrations were quantified by using Human E-Cadherin Quantikine ELISA kit (Invitrogen). The Human E-Cadherin EIA kit protocol was followed as described. Briefly, 100 μL of all sample or standard were added to appropriate wells prior to incubating the microtiter plate for 2 h at 37°C. The sample solutions were removed and each well was washed with 3 x 350 μL 1x Wash Buffer. Between each wash, the plate was emptied out and tapped vigorously onto paper towel, especially after the last washing. 100 μL of Antibody-HRP Conjugate Solution was added into each well followed by a 1 h incubation at 37°C. Sample solutions were removed and wells were washed 4 times as described above. 100 μL of Substrate Solution were added into each wells followed by a further incubation at 30°C for 30 min. 100 μL of Stop Solution was then added to each well before recording the absorbance at OD = 450 nm.

### Quantitative real-time PCR (qRT-PCR)

#### RNA extraction

RNA was either extracted from cell pellets that had previously been stored at −80°C, or from cell slurry. For lysis, 350μL RLT buffer supplemented with 1% pure β-mercaptoethanol were added and the cell slurry was transferred onto a QIAshredder spin column (Qiagen). Subsequently, cell lysates were spun on the QIashredder column and RNA extracted using the RNeasy kit (Qiagen) according to the manufacturer’s protocol. An optional on-column DNase treatment for 15 minutes was performed. RNA yields were measured by loading 1.5μL of extracted RNA onto the NanoDrop™ 2000c spectrophotometer. The optical density of each sample was subsequently verified for its purity (A 260/280 ratio = 2).

### cDNA production by reverse transcription (RT)

cDNA was produced by using either the Ambion RETROScript™ Kit (Invitrogen) or the High Capacity cDNA Reverse Transcription Kit (Applied Biosystems), and in both cases a total of 1μg RNA/sample was loaded for the cDNA reaction. For the TaqMan hPSC Scorecard™ Panel, cDNA was produced using the High Capacity cDNA Reverse Transcription Kit according to the manufacturer’s protocol.

### Quantitative real-time PCR (qRT-PCR)

qRT-PCR was either performed using a SYBR Green-(Applied Biosystems) or a TaqMan-based approach. To this end, the previously generated cDNA was diluted and prepared according to tables below. The primers for PKC isoforms and E-cadherin were adopted from Awadelkarim *et al.^iv^* and Labernadie *et al.^v^* and ordered from Sigma. Prior to all experiments, the efficiencies of the endogenous control (TBP) and the primer pairs were tested (Fig. S11).

**Table 1:** SYBR Green sample preparation

|  | Primer mix<br>(5 µL each) | 2 x SYBR<br>Select MM | Nuclease-free<br>H<sub>2</sub>O | Total volume |
| --- | --- | --- | --- | --- |
| ***SYBR Green reaction*** |  |  |  |  |
| Volume/sample + | 0.5 µL | 12.5 µL | 10 µL | 23 µL |
| 1.8 µL template cDNA (20 µL cDNA reaction diluted 1:10 in H<sub>2</sub>O) |  |  |  | 25 µL |

**Table 2:**
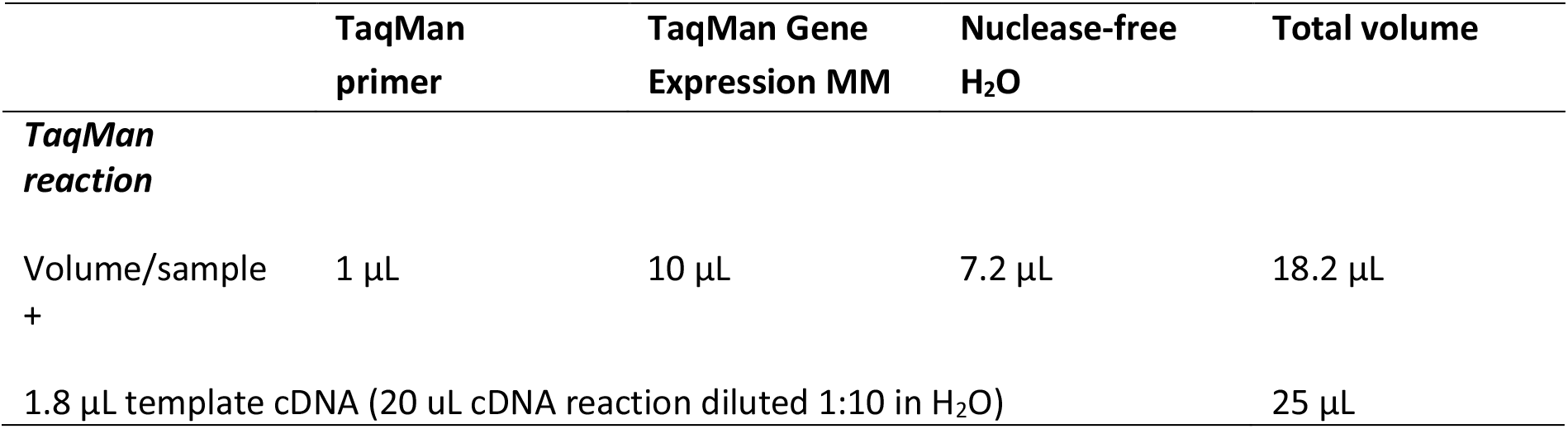
TaqMan sample preparation

The total volume of master mix and cDNA was loaded onto a MicroAmp™ Optical 96-Well Reaction Plate (Applied Biosystems), and the qRT-PCR reaction run on the StepOnePLus™ Real-Time PCR System (Applied Biosystems). Analysis of relative gene expression levels was performed according to the ΔΔCT approach^vi^.

### Enzyme-linked Immunosorbent Assay for PKC activity

PKC activity was tested using PKC kinase activity assay (Abcam) according to the manufacturer’s protocol. Brielfy, cells treated for 1 passage (5 days) with 0.1% of DMSO or 1 μM of OXS8360 were incubated with lysis buffer (20 mM MOPS, 50 mMβ-glycerolphosphate, 50 mM sodium fluoride, 1 mM sodium orthovanadate, 5 mM EGTA, 2 mM EDTA, 1% NP40, 1 mM DTT, 1 mM benzamidine, 1 mM PMSF,10μg/ml leupeptin and aprotinin) for 10 min on ice and then centrifuged at 13000 rpm for 15 min. 0.3 μg of protein from cell lysates diluted in 30μl of Kinase Assay Dilution Buffer were added to the pre-soaked wells of the PKC substrate microtiter plate. Standard diluted in 30μl of Kinase Assay Dilution Buffer and 30 μl of Kinase Assay Dilution Buffer (blank) were also added in appropriate wells. The kinase reaction was initiated by adding 10 μl of ATP to each well and the samples were incubated for 90 min at 30°C with gentle shaking after 20 min. The reaction was stopped by removing the contents of each well. Samples, excluding blank ones, were incubated with 40 μl of Phosphospecific substrate antibody for 1 h at room temperature with gentle shaking every 20 min. Wells were washed four times with Wash Buffer and then incubated with 40 μL of diluted anti-rabbit IgG-HRP antibody for 30 min at room temperature, with gentle shaking every 10 min. Subsequently, all wells were washed four times with Wash Buffer. In order to detect PKC activity, 60 μL of TMB substrate was added to each well and the plate was incubated at room temperature for 60 min and then stopped by addition of 20 μL of Stop Solution. The PKC activity was analyzed by measuring the absorbance at OD = 450 nm.

## Acknowledgments

Financial support: The Wellcome Trust WTISSF121302 and the Oxford Martin School LC0910-004 (James Martin Stem Cell Facility, Oxford, S.A.C.). The research leading to these results has received support from the Innovative Medicines Initiative Joint Undertaking under grant agreement n° 115439, resources of which are composed of financial contribution from the European Union’s Seventh Framework Programme (FP7/2007– 2013) and EFPIA companies’ in kind contribution. This publication reflects only the author’s views and neither the IMI JU nor EFPIA nor the European Commission are liable for any use that may be made of the information contained therein. This work was also supported by funding from Oxstem Limited. We thank the High-Throughput Genomics Group at the Wellcome Trust Centre for Human Genetics, Oxford (Funded by Wellcome Trust grant reference 090532/Z/09/Z and MRC Hub grant G0900747 91070) for the generation of Illumina SNP data.

## Author contributions

Conceptualization, S.A.C., A.J.R., W.S.J., L.S., G.L. and S.G.D.; Formal analysis, L.S., M.Sc, G.L., M.O., L.G.-P., E.C., C.J.R.B. and M.St.; Funding acquisition, S.A.C., A.J.R., W.S.J. and S.G.D.; Investigation, L.S., M.Sc., G.L., M.O., E.C., C.J.R.B. and M.St.; Methodology, L.S., M.Sc., G.L., M.O., L.G.-P., E.C., C.J.R.B. and M.St.; Resources, S.A.C., A.J.R., W.S.J. and L.W.S.; Supervision, S.A.C., W.S.J. and A.J.R.; Writing – original draft, L.S.; Writing – review and editing, S.A.C. and A.J.R.

