## Supplementary Information for "A chemical tool for improved culture of human pluripotent stem cells"

**Supporting information**

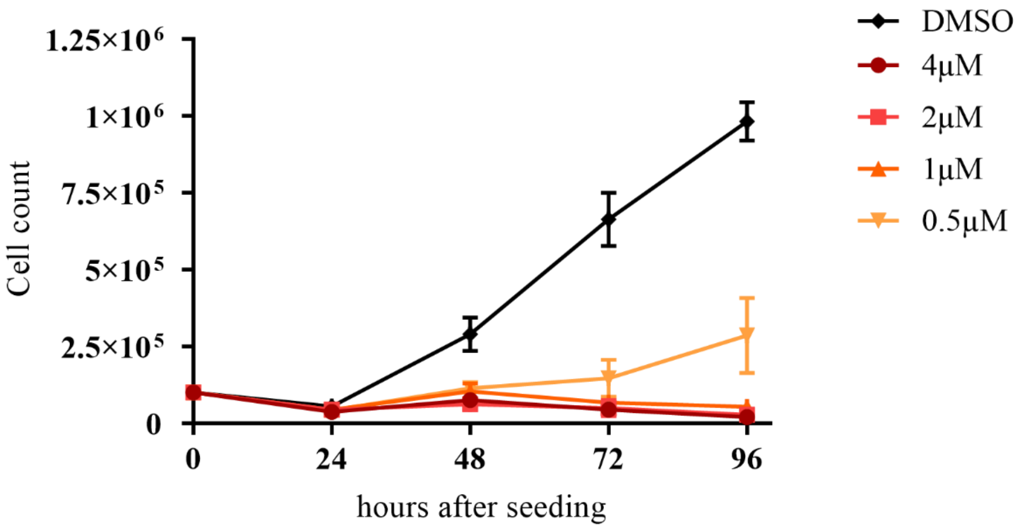


**Figure S1:** Growth curve of OX1-18 iPSC treated with (-)-Indolactam V. Mean of three separate wells ±SEM, for each time point.

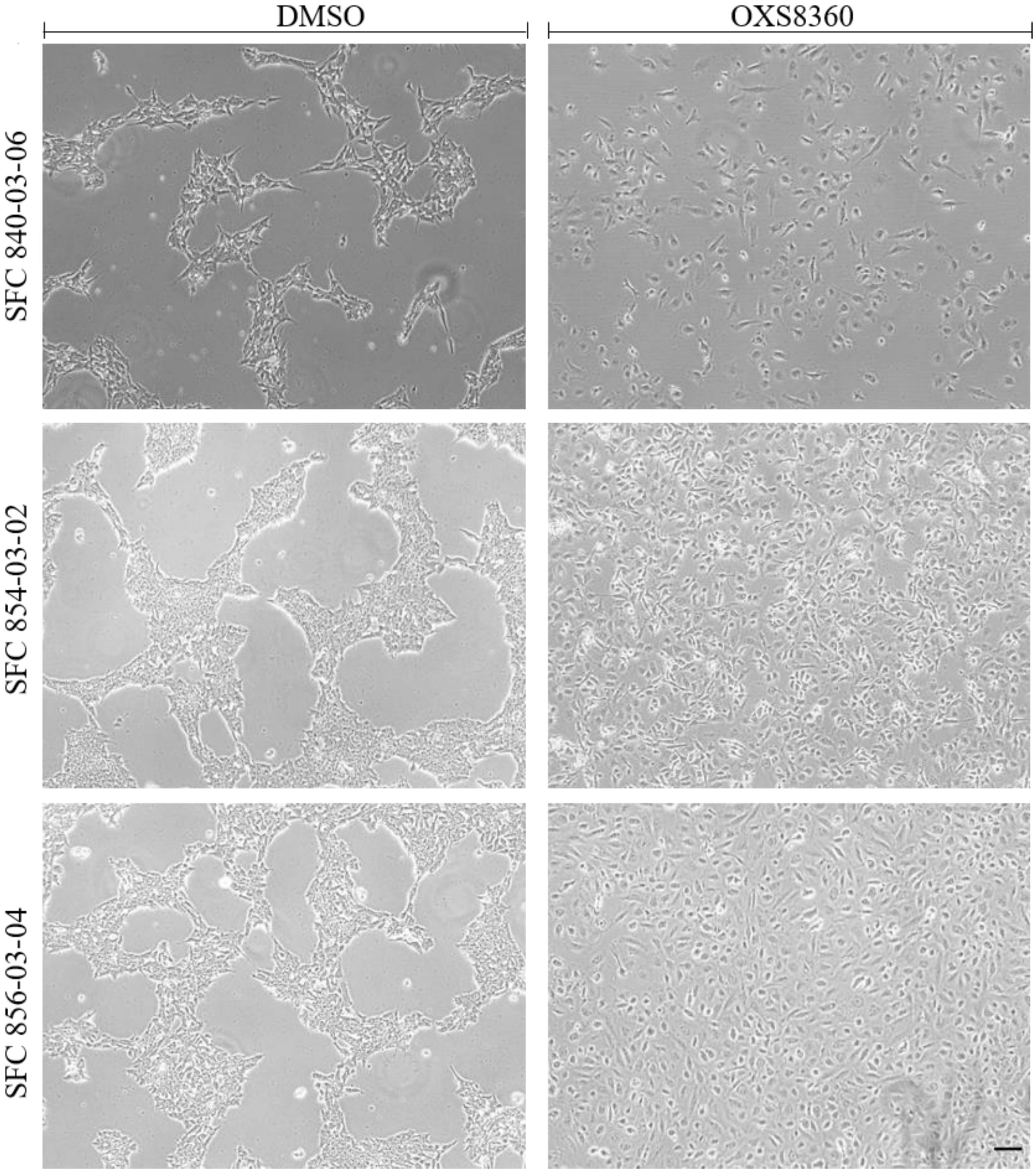


**Figure S2:** Phenotypic effect of OXS8360 on 3 genetically different iPSC lines.
SFC 840-03-06, SFC 854-03-02 and SFC 856-03-04 iPSC lines were treated with DMSO or OXS8360 for 48h. The cell-spreading effect can be observed in all 3 lines. Phase contrast microscopy. Scale bar = 100 µm.

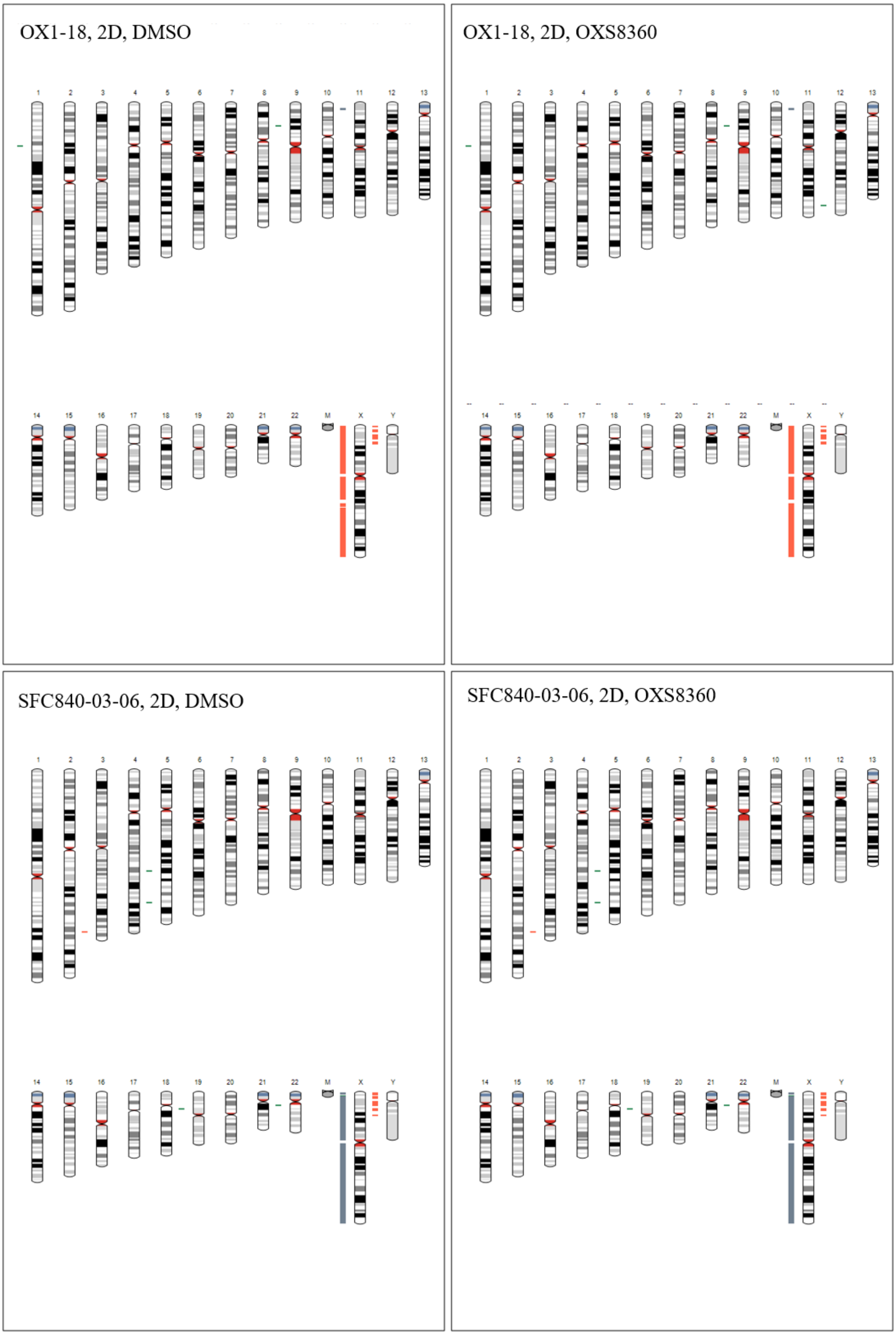
OX1-18, 2D, DMSO
OX1-18, 2D, OXS8360
SFC840-03-06, 2D, DMSO
SFC840-03-06, 2D, OXS8360

SFC854-03-02, 2D, DMSO

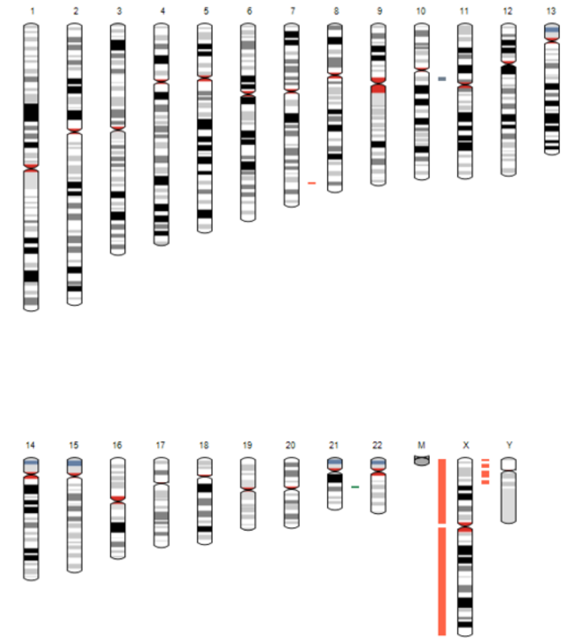

SFC854-03-02, 2D, OXS8360

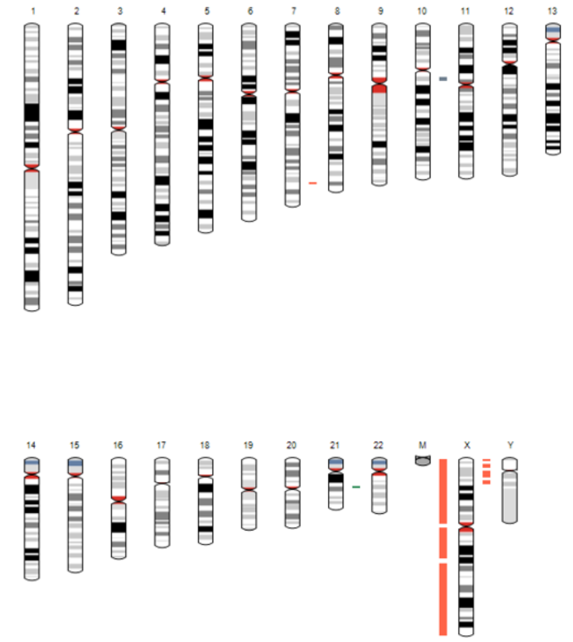

SFC856-03-04, 2D, DMSO

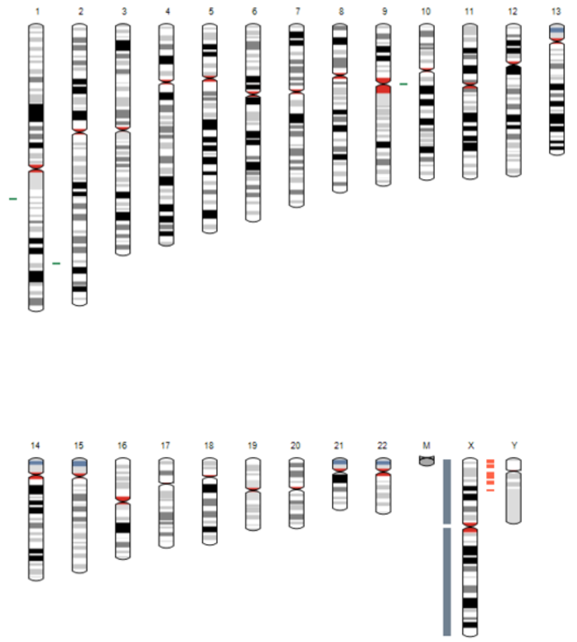

SFC856-03-04, 2D, OXS8360

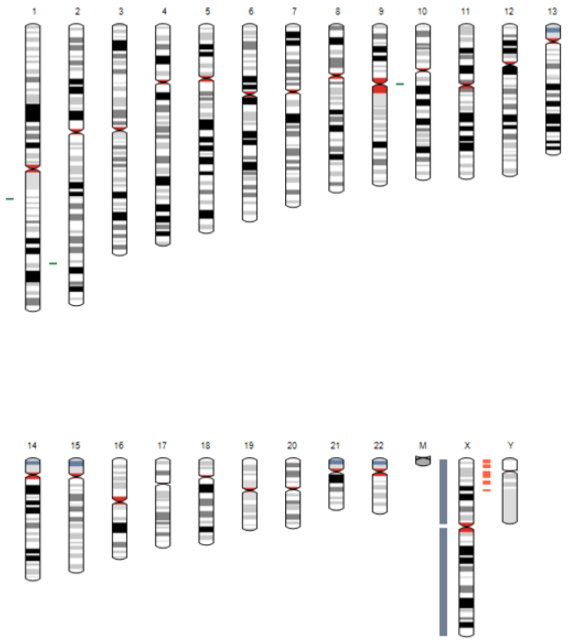

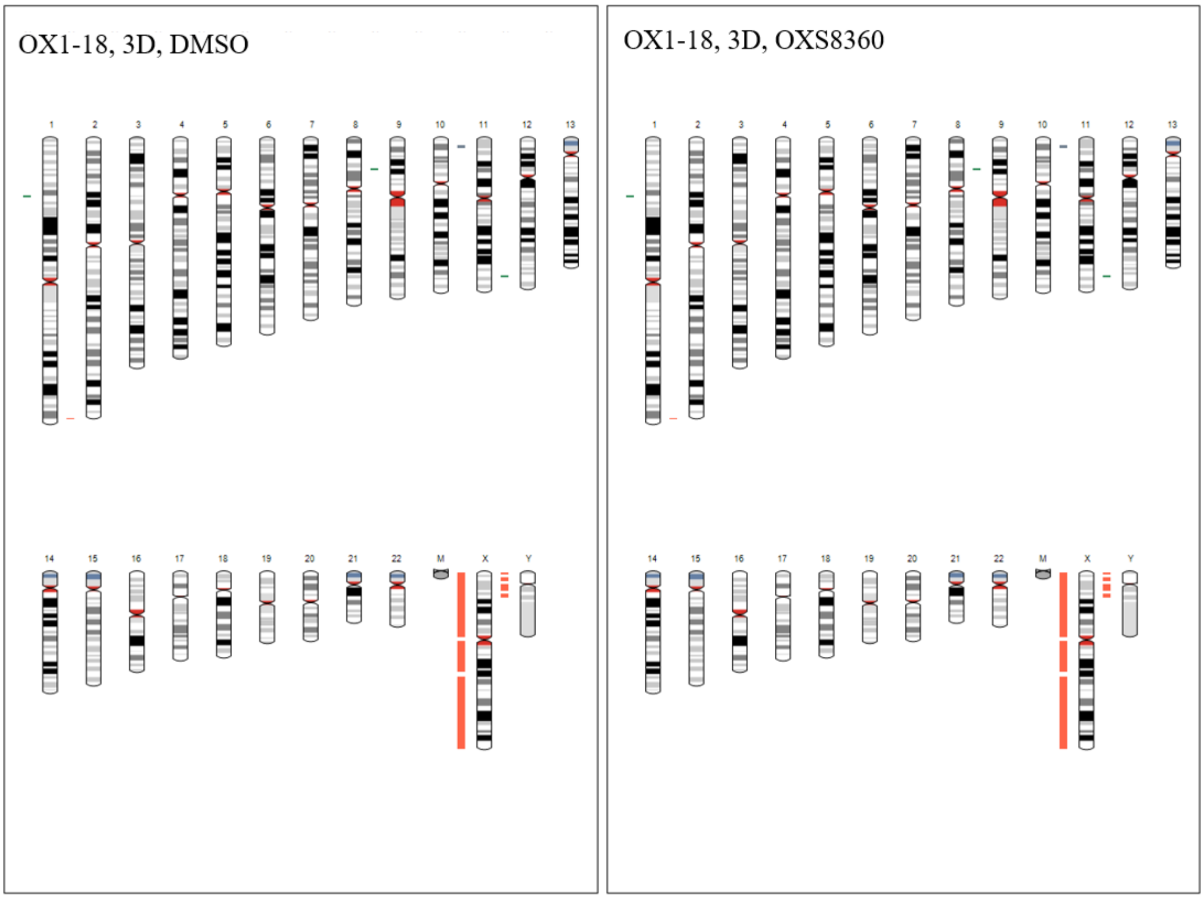


**Figure S3**: Genome integrity analysis of four iPSC clones (OX1-18, SFC840-03-06, SFC854-03-02, SFC856-03-04) grown in 2D or 3D suspension culture.

The analysis indicates a normal chromosome complement and no gross structural abnormalities in OXS8360-treated iPSC over 10 passages when compared to the control (DMSO). Red bar adjacent to the chromosome indicates loss or single copy, green indicates gain of copy, grey indicates loss of heterozygosity on autosomes, or two copies of X chromosome (i.e. female lines)

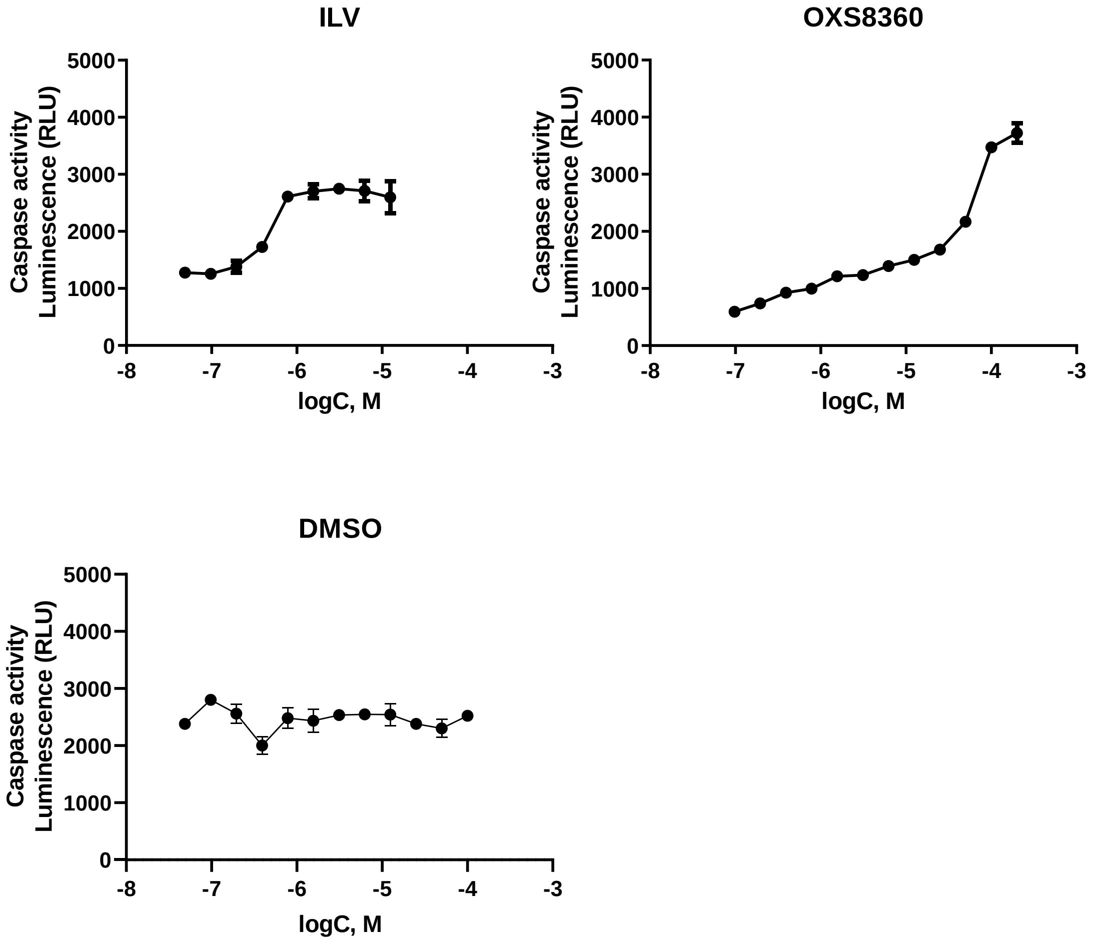


**Figure S4:** Caspase activity assay.

$IC_{50}$ values for (-)-ILV and OXS8360 for apoptosis were determined by monitoring caspase activity by using Caspase-Glo 3/7 assay kit after OX1-18 iPSC were treated with DMSO, (-)-ILV and OXS8360 for 24h. $IC_{50}$ values were calculated using Graphpad Prism and determined as 5 µM and 94 µM for (-)-ILV and OXS8360 respectively. Mean±SEM, n=3.

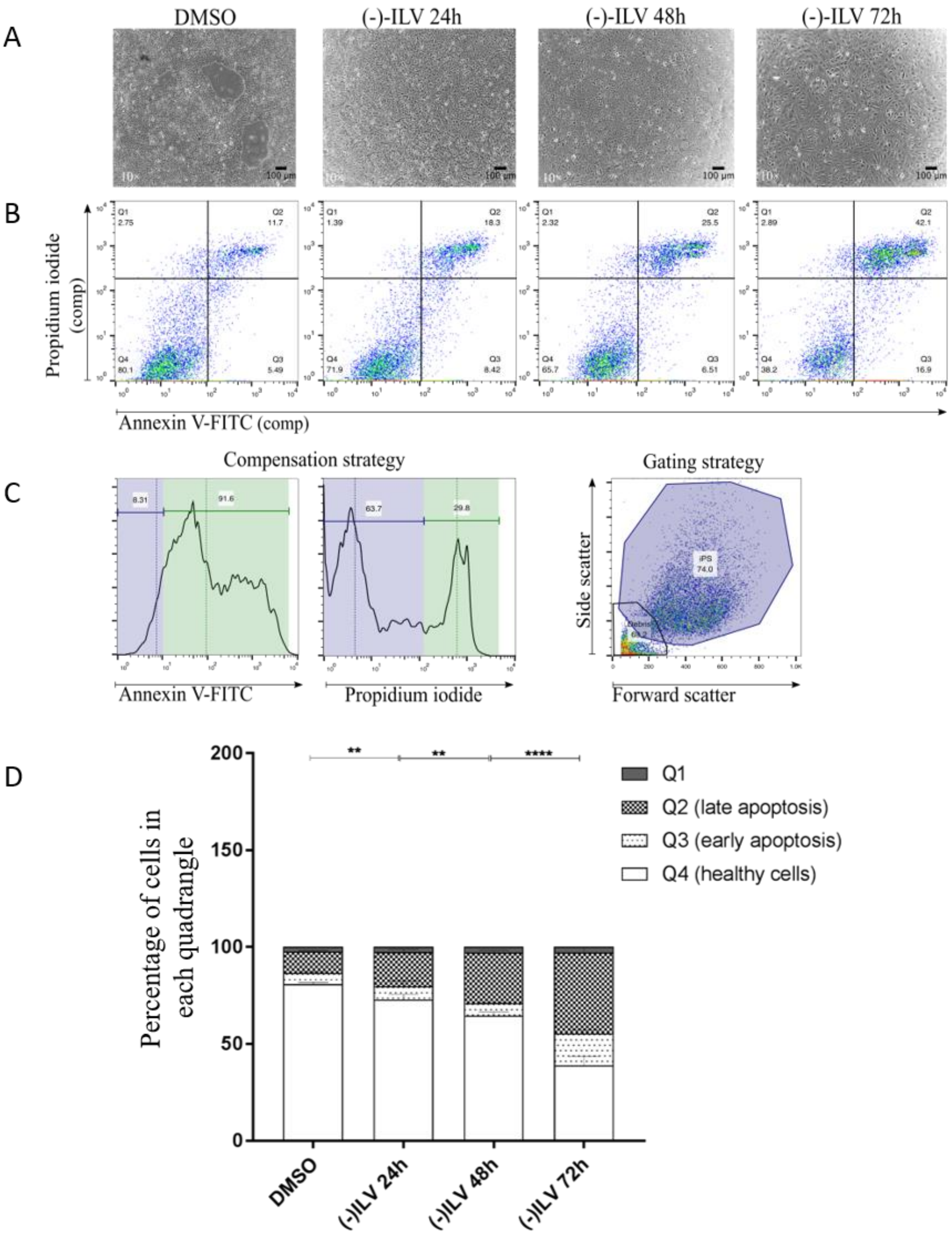


**Figure S5:** Analysing the cytotoxic effects of (-)-ILV using an Annexin-PI assay. **(A)** Phase contrast microscopy of OX1-18 iPSC treated with (-)-ILV. The phenotypic effect can be observed in all treated conditions, with an apparent decrease of cell density at 72h. Scale bar = 100 µm **(B)** All conditions were stained with Annexin-V and PI and analysed using flow cytometry. Double negative populations (Q4) are healthy, single positive populations for Annexin-V have entered early apoptosis (Q3), double positive populations have entered late apoptosis and undergo necrosis (Q2). **(C)** Populations in (B) were previously compensated in the PI channel to avoid false positive signals due to spill over from the FITC-Annexin-V channel. During the gating process, debris is excluded. **(D)** Mean ± SEM of each population in Q1-Q4 is plotted in stacked bars. DMSO (2.34±0.89; 11.13±0.30; 5.56±0.45; 80.97±0.47; n=3); (-)-ILV 24h (2.74±0.99; 17.47±0.88; 6.96±1.62; 72.83±1.57; n=3); (-)-ILV 48h (2.82±0.54; 26.07±0.67; 6.61±0.11; 64.50±1.15; n=3); (-)-ILV 72h (2.97±0.09; 41.53±1.67; 16.57±1.05; 38.93±2.68; n=3). Differences in Q1 for all conditions and in Q3 for DMSO, (-)-ILV 24h and (-)-ILV 48h are not significant. Two-way ANOVA; Holm-Sidak's multiple comparison test.

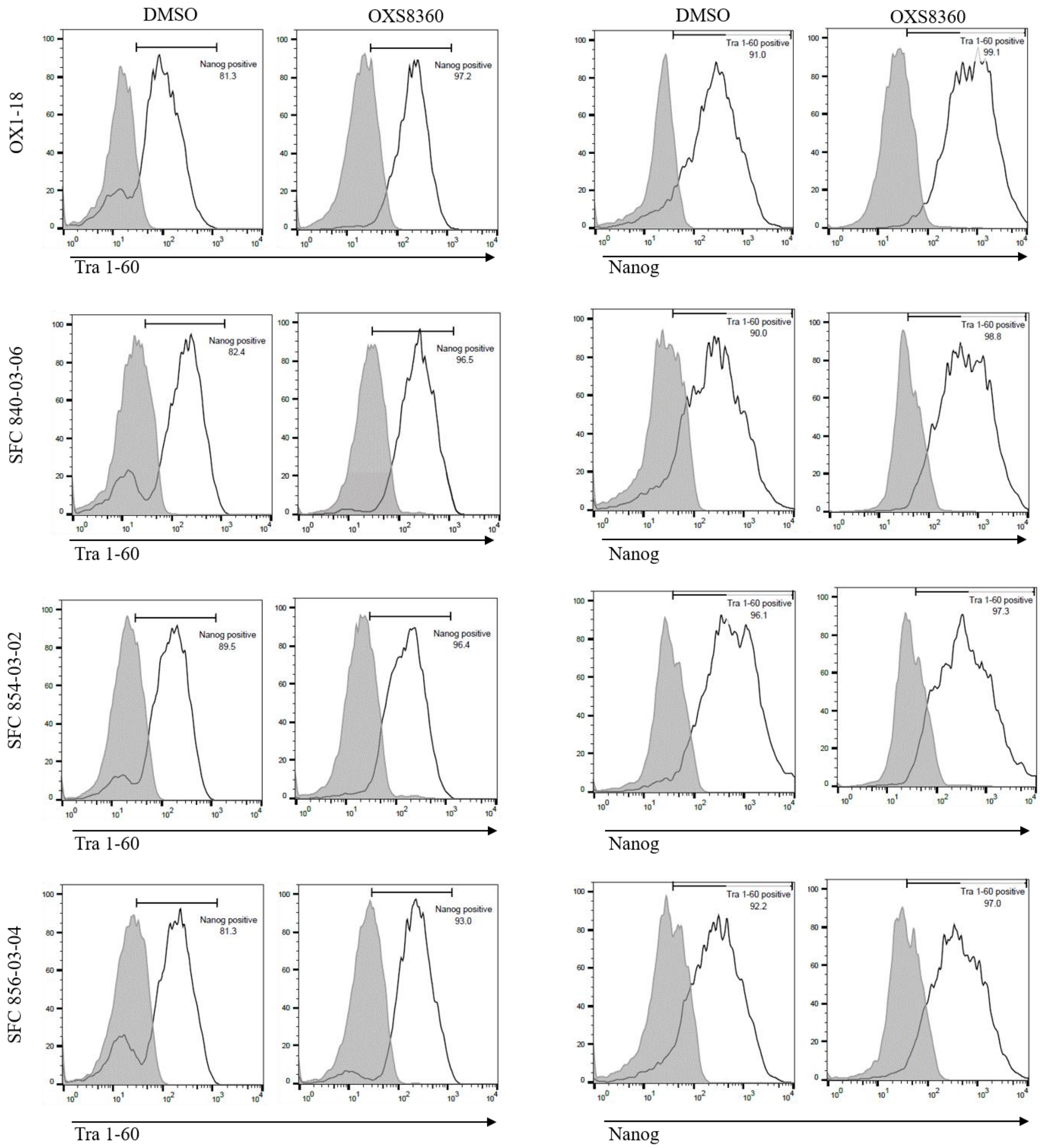


**Figure S6**: Expression of pluripotency markers after prolonged compound treatment.
Flow cytometry for pluripotency markers Tra1-60 and Nanog for iPSC lines OX1-18, SFC840-03-06, SFC854-03-02, SFC856-03-04 treated with DMSO or OXS8360 over 10 passages.

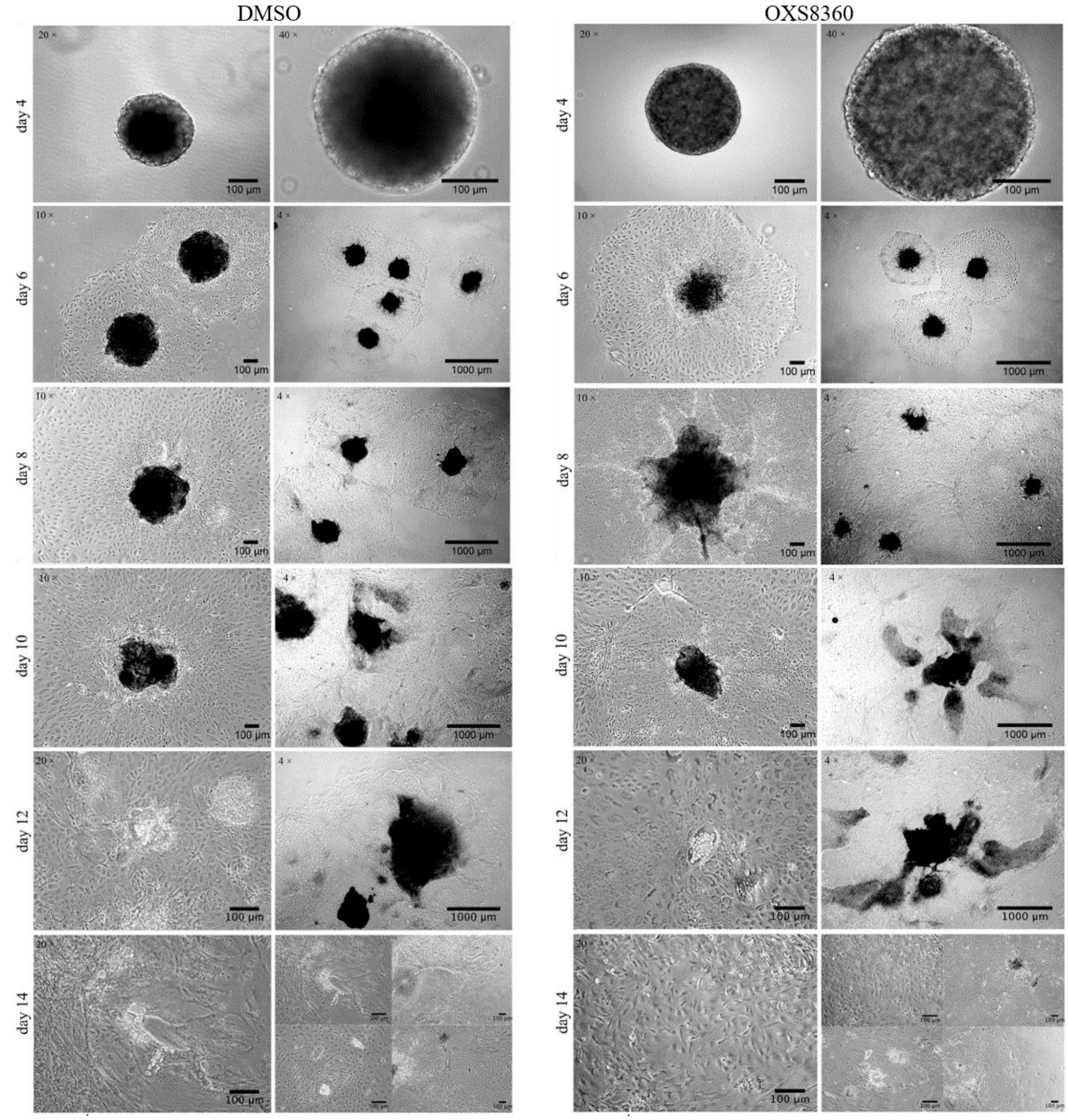


**Figure S7**: Undirected differentiation of iPSC OX1-18 treated with DMSO or OXS8360 prior to differentiation. Day 4, spin-EBs transferred from Aggrewells™ to a Geltrex™ coated dish. Day 6, the EBs have attached and the cells start to grow out from the centre of the EBs. Day 12 and 14, cell density has increased and the cell populations become increasingly heterogenous. Phase contrast microscopy. Scale bars are as indicated.

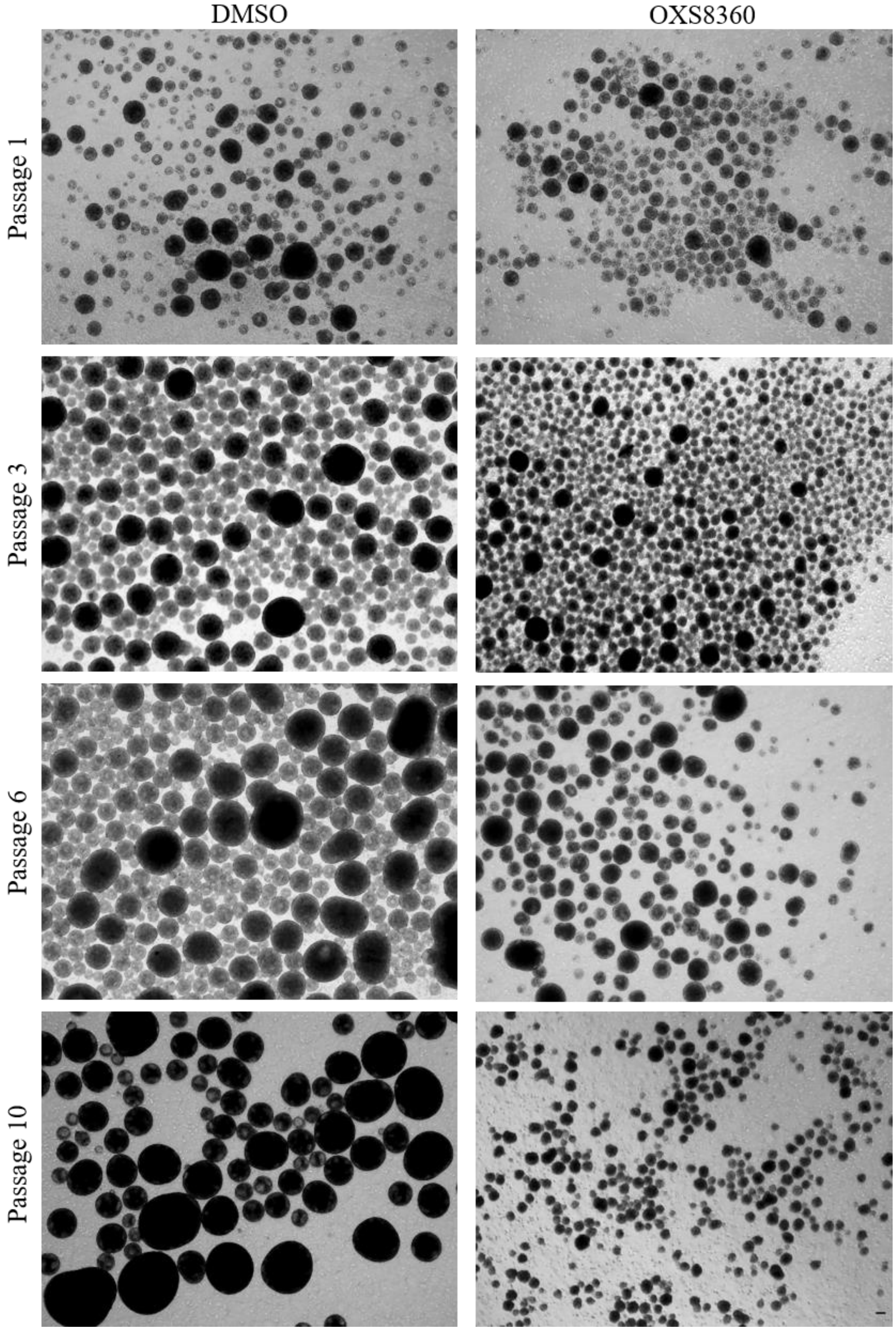


**Figure S8:** Effect of OXS8360 on hPSC aggregate size when grown in suspension culture over multiple passages.
0.5x$10^6$ cells were seeded in each well and treated with DMSO or OXS8360 at 2 µM. Images were taken every 4 days, just before passaging. Phase contrast microscopy. Scale bar = 100 µm.

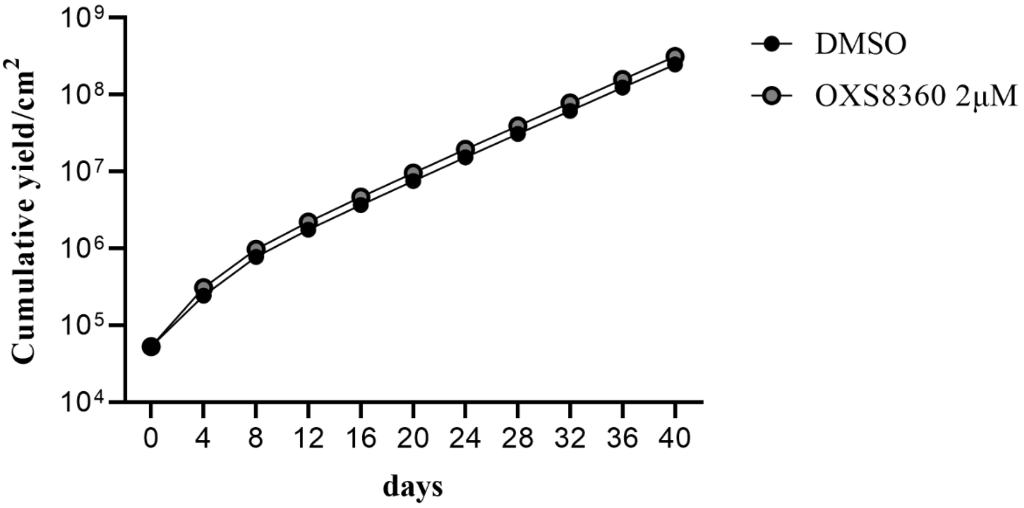


**Figure S9:** Yields of OX1-18 iPSC obtained from 3D static suspension culture.
iPSC were sub-cultured every 4 days at a density of $0.5 \times 10^6$ cells/well for 40 days (10 passages). At each time point cells were counted in triplicate. **(A)** The cumulative yield of DMSO and OXS8360-treated cells, normalised to tissue culture plastic area, over 40 days

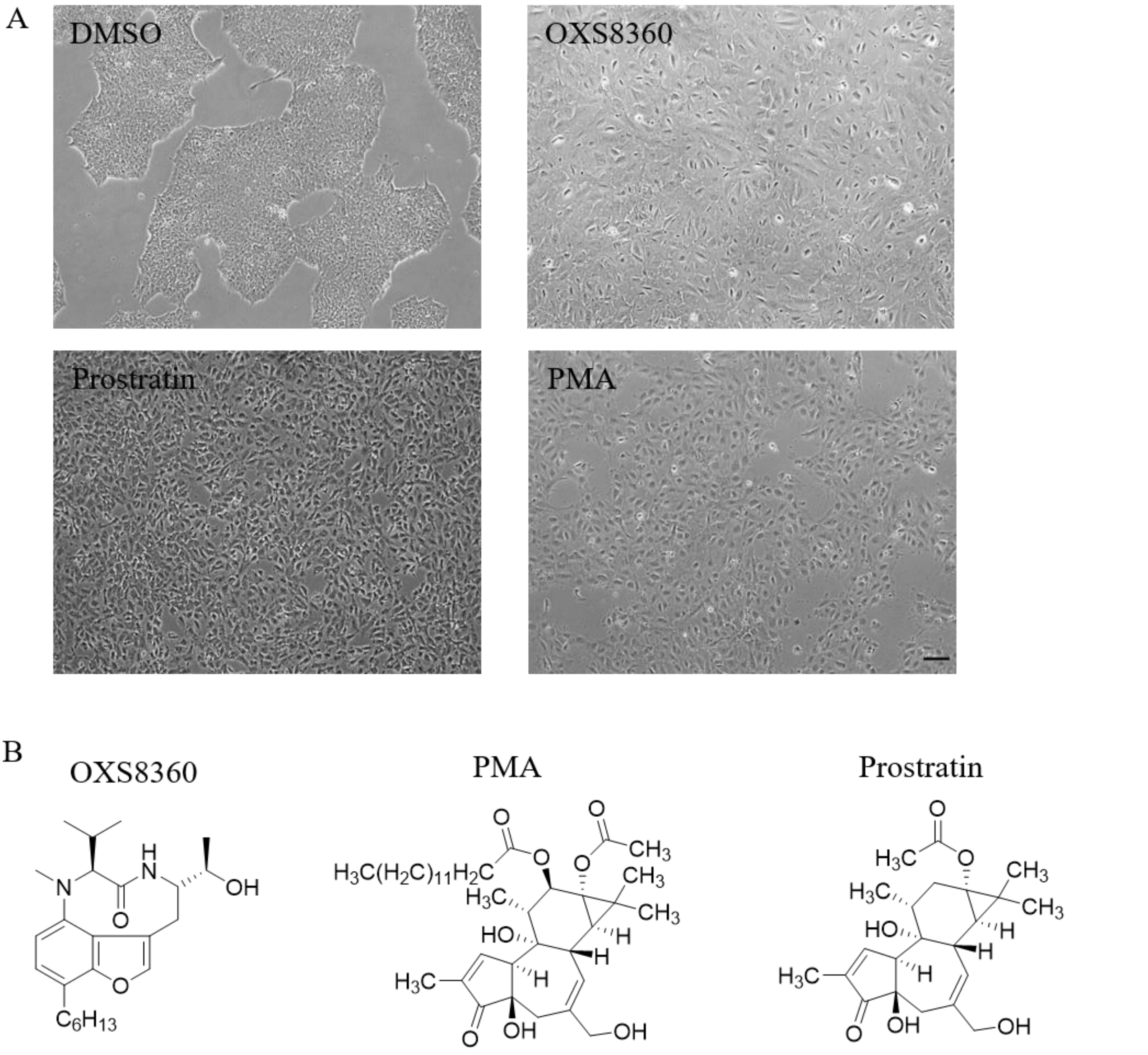


**Figure S10:** Phenotypic effect of PKC activators on OX1-18 iPSC. **(A)** OX1-18 iPSC treated with 1 µM of commercially available PKC activators (PMA and Prostratin) versus OXS8360. Cell spreading can be seen in all conditions. **(B)** Chemical structure of PKC activators. Phase contrast microscopy. Scale bar = 100 µm.

**Table S1**: iPSC lines used in this study

| StemBANCC/ OPDC of iPSC clone | sex | age | reprogramming method | publication | deposited in repository |
|---|---|---|---|---|---|
| OX1-18 | M | 36 | Yamanaka retrovirus | PMID:23951090 | EBiSC |
| SFC840-03-06 | F | 67 | Cytotune 1 | PMID:28827786 | EBiSC |
| SFC854-03-02 | M | 72 | Cytotune 2 | PMID:28827786 | EBiSC |
| SFC856-03-04 | F | 78 | Cytotune 2 | PMID:28827786 | EBiSC |

**Table S2**: ScoreCard assay Ct values

| Gene | Category | DMSO undiff. (1) | OXS8360 undiff. (2) | DMSO diff. (3) | OXS8360 diff. (4) |
|---|---|---|---|---|---|
| ACTB | Controls | 19.16 | 19.19 | 19.23 | 18.98 |
| ACTB | Controls | 24.23 | 22.85 | 22.97 | 22.39 |
| ACTB | Controls | 23.72 | 22.96 | 22.70 | 21.80 |
| ACTB | Controls | 19.56 | 19.56 | 19.57 | 18.99 |
| CTCF | Controls | 25.94 | 25.96 | 26.37 | 26.28 |
| EP300 | Controls | 25.85 | 25.94 | 26.58 | 26.30 |
| SMAD1 | Controls | 27.78 | 27.97 | 27.66 | 27.75 |
| CDH9 | Ectoderm | 33.41 | 32.42 | 32.90 | 32.92 |
| COL2A1 | Ectoderm | 30.35 | 29.92 | 25.95 | 27.21 |
| DMBX1 | Ectoderm | 32.97 | 30.87 | 31.26 | 31.06 |
| DRD4 | Ectoderm | 28.41 | 27.91 | 26.97 | 28.57 |
| EN1 | Ectoderm | 36.95 | 35.97 | 31.10 | 31.74 |
| LMX1A | Ectoderm | 35.95 | 34.71 | 28.41 | 31.49 |
| MAP2 | Ectoderm | 29.27 | 28.83 | 27.96 | 29.76 |
| MYO3B | Ectoderm | 37.02 | 35.91 | 31.57 | 32.42 |
| NOS2 | Ectoderm | 30.83 | 30.97 | 27.86 | 29.43 |
| NR2F1/NR2F2 | Ectoderm | 36.96 | 34.30 | 24.14 | 26.39 |
| NR2F2 | Ectoderm | 36.96 | 35.25 | 25.01 | 24.73 |
| OLFM3 | Ectoderm | 33.95 | 33.54 | 30.16 | 32.80 |
| PAPLN | Ectoderm | 28.79 | 27.98 | 31.11 | 31.48 |
| PAX3 | Ectoderm | 36.99 | 40.00 | 25.88 | 28.34 |
| PAX6 | Ectoderm | 35.84 | 34.23 | 26.98 | 29.86 |
| POU4F1 | Ectoderm | 34.87 | 35.07 | 28.98 | 30.98 |
| PRKCA | Ectoderm | 27.19 | 26.92 | 28.44 | 28.69 |
| SDC2 | Ectoderm | 25.01 | 25.47 | 25.74 | 25.91 |
| SOX1 | Ectoderm | 36.93 | 35.90 | 33.08 | 36.96 |
| TRPM8 | Ectoderm | 36.48 | 35.97 | 34.36 | 35.84 |
| WNT1 | Ectoderm | 37.01 | 35.77 | 25.83 | 27.67 |
| ZBTB16 | Ectoderm | 33.83 | 33.97 | 27.74 | 30.43 |
| AFP | Endoderm | 37.05 | 40.00 | 24.24 | 22.16 |
| CABP7 | Endoderm | 33.65 | 33.65 | 32.07 | 32.99 |
| CDH20 | Endoderm | 34.44 | 34.92 | 28.96 | 30.57 |
| CLDN1 | Endoderm | 29.88 | 29.91 | 27.96 | 28.30 |
| CPLX2 | Endoderm | 31.86 | 31.66 | 29.92 | 31.41 |
| ELAVL3 | Endoderm | 31.59 | 31.98 | 29.65 | 30.77 |
| EOMES | Endoderm | 34.92 | 34.89 | 27.94 | 28.54 |
| FOXA1 | Endoderm | 35.69 | 35.99 | 29.30 | 29.62 |
| FOXA2 | Endoderm | 35.78 | 37.01 | 27.09 | 27.90 |
| FOXP2 | Endoderm | 35.76 | 35.94 | 30.30 | 31.01 |
| GATA4 | Endoderm | 33.56 | 34.29 | 26.95 | 26.38 |
| GATA6 | Endoderm | 35.81 | 35.63 | 26.30 | 25.92 |
| HHEX | Endoderm | 30.19 | 29.64 | 29.99 | 29.98 |
| HMP19 | Endoderm | 34.72 | 34.95 | 33.81 | 34.60 |
| HNF1B | Endoderm | 33.85 | 33.77 | 29.90 | 29.96 |
| HNF4A | Endoderm | 36.99 | 35.41 | 25.44 | 25.02 |
| KLF5 | Endoderm | 32.64 | 32.61 | 28.94 | 28.47 |
| LEFTY1 | Endoderm | 33.95 | 33.77 | 33.81 | 34.39 |
| LEFTY2 | Endoderm | 33.92 | 34.49 | 32.62 | 33.00 |
| NODAL | Endoderm | 30.95 | 31.87 | 27.95 | 28.94 |
| PHOX2B | Endoderm | 40.00 | 40.00 | 34.60 | 35.99 |

**Table S2**: ScoreCard assay Ct values (continued)

| Gene | Category | DMSO undiff. (1) | OXS8360 undiff. (2) | DMSO diff. (3) | OXS8360 diff. (4) |
|---|---|---|---|---|---|
| FGF4 | Mesendoderm | 32.20 | 32.49 | 29.07 | 29.96 |
| GDF3 | Mesendoderm | 29.43 | 29.94 | 27.50 | 28.98 |
| NPPB | Mesendoderm | 36.97 | 35.53 | 25.24 | 24.47 |
| NR5A2 | Mesendoderm | 28.82 | 28.81 | 31.66 | 31.96 |
| PTHLH | Mesendoderm | 35.66 | 36.97 | 29.70 | 27.40 |
| T | Mesendoderm | 35.99 | 35.50 | 30.08 | 30.07 |
| ABCA4 | Mesoderm | 35.93 | 35.62 | 29.12 | 29.45 |
| ALOX15 | Mesoderm | 30.67 | 30.53 | 26.94 | 25.70 |
| BMP10 | Mesoderm | 40.00 | 40.00 | 32.98 | 32.92 |
| CDH5 | Mesoderm | 34.83 | 35.01 | 26.87 | 26.01 |
| CDX2 | Mesoderm | 35.38 | 35.99 | 27.38 | 28.00 |
| COLEC10 | Mesoderm | 33.67 | 33.73 | 31.95 | 29.88 |
| ESM1 | Mesoderm | 40.00 | 40.00 | 29.95 | 29.96 |
| FCN3 | Mesoderm | 32.89 | 32.63 | 31.95 | 31.49 |
| FOXF1 | Mesoderm | 36.42 | 36.95 | 27.57 | 26.74 |
| HAND1 | Mesoderm | 36.43 | 36.96 | 22.91 | 22.47 |
| HAND2 | Mesoderm | 36.71 | 35.92 | 25.97 | 25.04 |
| HEY1 | Mesoderm | 32.24 | 31.94 | 26.68 | 27.42 |
| HOPX | Mesoderm | 37.21 | 35.56 | 23.10 | 22.51 |
| IL6ST | Mesoderm | 29.88 | 29.97 | 25.99 | 24.87 |
| NKX2-5 | Mesoderm | 35.56 | 35.95 | 32.43 | 31.67 |
| ODAM | Mesoderm | 36.78 | 37.10 | 27.60 | 28.33 |
| PDGFRA | Mesoderm | 32.76 | 32.51 | 26.96 | 26.73 |
| PLVAP | Mesoderm | 33.71 | 33.96 | 28.14 | 26.98 |
| RGS4 | Mesoderm | 34.53 | 34.94 | 26.93 | 26.88 |
| SNAI2 | Mesoderm | 32.83 | 33.24 | 25.95 | 25.28 |
| TBX3 | Mesoderm | 33.03 | 32.57 | 24.19 | 23.94 |
| TM4SF1 | Mesoderm | 34.40 | 34.86 | 28.99 | 27.87 |
| CD44 | Other | 40.00 | 40.00 | 40.00 | 40.00 |
| JARID2 | Other | 23.90 | 23.59 | 25.38 | 25.97 |
| MYC | Other | 25.96 | 25.49 | 26.96 | 25.89 |
| SEV | Other | 40.00 | 40.00 | 40.00 | 40.00 |
| CXCL5 | Self-renewal | 25.96 | 26.30 | 30.56 | 31.29 |
| DNMT3B | Self-renewal | 20.68 | 20.87 | 22.95 | 23.89 |
| HESX1 | Self-renewal | 29.39 | 29.33 | 31.75 | 31.94 |
| IDO1 | Self-renewal | 25.93 | 25.93 | 30.61 | 31.62 |
| LCK | Self-renewal | 25.73 | 25.95 | 29.68 | 30.65 |
| NANOG | Self-renewal | 24.93 | 24.92 | 26.06 | 27.41 |
| POU5F1 | Self-renewal | 28.47 | 28.60 | 31.21 | 31.95 |
| SOX2 | Self-renewal | 23.19 | 22.95 | 24.57 | 26.78 |
| TRIM22 | Self-renewal | 25.92 | 26.55 | 28.58 | 28.18 |
